# MicroRNA spatial profiling for assessing drug efficacy in *BRCA1*-related triple-negative breast tumors

**DOI:** 10.64898/2026.01.12.698824

**Authors:** Omar N. Mohd, Lin Wang, Brian R. Sardella, David Jou, Gerburg M. Wulf, Frank J. Slack, Yujing J. Heng, Patrick S. Doyle

## Abstract

*BRCA1/2*-mutated breast cancers exhibit homologous recombination deficiency (HRD) and are initially sensitive to poly(ADP-ribose) polymerase (PARP) inhibitors, but 40-70% of patients develop resistance, creating a need for predictive biomarkers that capture treatment-associated spatial heterogeneity. Using the *K14-Cre Brca1^f/f^Trp53^f/f^*model with tumors that acquired PARP inhibitor resistance, we evaluated PARP inhibitor combinations with either PI3K inhibition or Poly(I:C) *in vivo*. To determine how treatment altered tumor spatial microRNA (miRNA) profiles, we applied our hydrogel-based, nanoliter well array *in situ* miRNA assay to quantify and spatially profile miRNAs on FFPE sections from tumors treated for 10 days and developed spatial miRNA analysis frameworks integrating latent Dirichlet allocation (LDA) and principal component analysis (PCA). We also incorporated immune architecture using Structural Similarity Index Measure (SSIM) maps to assess local structural similarity of immune infiltration and miRNA topics. Both combinations improved antitumor activity compared to PARP inhibition alone in olaparib-sensitive tumors. The resulting spatial miRNA topics stratified early tumors according to established PARP inhibitor sensitivity or resistance and distinguished their treatment regimens, while SSIM analysis revealed local structural similarity of immune infiltration and miRNA topics. This integrative spatial miRNA assay and analysis identified a let-7a-dominant topic associated with PARP inhibitor resistance, indicating that spatial miRNA profiling may inform therapeutic stratification in *BRCA1/2*-related breast cancers.

## Introduction

Breast tumors with a germline *BRCA1/2* mutation have a defective homologous recombination DNA double-strand break repair pathway.[1] Poly(ADP-ribose) polymerase (PARP) inhibitors are FDA-approved to treat advanced *BRCA1/2*-related breast cancers.[2] PARP inhibitors work by stalling and collapsing replication forks thereby stopping DNA repair and eventual cell death.[3,4] Nevertheless, about 40% to 70% of patients fail to respond or will develop resistance to PARP inhibitors over time.[5–7] The high rate of resistance has driven efforts to develop combination therapies that enhance the effectiveness of PARP inhibitors, as well as to identify biomarkers to predict which patients are most likely to benefit.

MicroRNAs (miRNAs) are small (∼20 nucleotides), non-coding RNAs that have emerged as useful biomarkers for the detection of early cancers, including breast cancer.[8–12] As such, miRNAs may provide informative molecular readouts of PARP inhibitor response and resistance in breast tumors. However, miRNA measurements are often performed in bulk tissue, which obscures intratumoral heterogeneity and removes spatial information.[13] We previously showed that integrated analysis of expression and spatial localization of seven miRNAs in endpoint murine breast tumors from untreated recipient mice discriminated PARP inhibitor-sensitive from -resistant tumors.[14] These findings suggest that spatially resolved miRNA measurements can capture treatment-associated tumor states that are not readily accessible from bulk assays.

One strategy to overcome PARP inhibitor resistance involves co-targeting PARP and the PI3-kinase (PI3K) pathway. PI3K inhibitors can suppress the growth of tumors with *PIK3CA* mutations,[15] and are also effective in tumors with wild-type *PIK3CA*.[16] In addition to their role in tumor proliferation, PI3K enzymes contribute to the DNA damage response: *PIK3CB* is required for double-strand break sensing by regulating NBN recruitment to damaged DNA.[17] Building on this mechanistic rationale, we and others have demonstrated that combining a PARP inhibitor with a PI3K inhibitor produces significant preclinical and clinical activity in BRCA1/2-related cancers.[18–21] We have also previously published that the effectiveness of PARP inhibition depends on the tumor immune microenvironment (TME).[22–24] Olaparib increased infiltration and activation of CD8^+^ T-cells [23] and tumor-associated macrophages.[22] Therefore, another potential combination therapy could be combining PARP with an immune stimulant such as polyinosinic:polycytidylic acid (Poly(I:C)). Poly(I:C) activates dendritic cells by binding to various surface receptors such as toll-like receptor 3 which in turn activates CD8^+^ T-cells.[25] These mechanistically distinct treatment conditions provide a biologically relevant setting in which to evaluate spatial molecular readouts of treatment response and resistance.

In this work, we used our previously described hydrogel-based, nanoliter well array *in situ* miRNA technology [14,26] to assess tumor spatial miRNA profiles in FFPE sections after 10 days of treatment, rather than at the treatment endpoint examined previously.[14] To interpret this high-dimensional spatial readout, we developed an analytical framework integrating latent Dirichlet allocation (LDA) and principal component analysis (PCA) to identify early, spatially resolved miRNA profiles associated with treatment response. We further developed a specialized, priming-based LDA analysis to improve differentiation between treatment regimens among PARP inhibitor-sensitive tumors. Finally, we evaluated whether incorporating intratumoral immune composition, by computing Structural Similarity Index Measure (SSIM)[27] maps between immune cell infiltration and miRNA topics, could enhance interpretation of the LDA-derived spatial topics by relating them to underlying cellular architecture and highlighting regions where immune patterns and miRNA distributions exhibit local structural similarity. This integrative spatial miRNA assay and analysis provides a framework for linking treatment-associated miRNA patterns to tumor and immune architecture and may inform treatment-response stratification in BRCA1/2-related breast cancers.

## Results

### Combination therapies produced promising therapeutic effects in PARP inhibitor-resistant tumors

We used the *K14-Cre Brca1^f/f^ Trp53^f/f^* mouse tumor model to investigate the effects of combining a PARP inhibitor, olaparib, with either a PI3K inhibitor (i.e., alpelisib) or Poly(I:C) (**Fig 1A**). This *BRCA1*-related triple-negative breast cancer (TNBC) model is known to be sensitive to PARP inhibition [28], as well as in combination with alpelisib.[18] Our current results using orthotopic xenotransplanted tumor fragments from this model were consistent with prior findings: olaparib significantly lengthened the median time taken to reach study endpoint of 20 mm compared to control (75-78 days versus 19-23 days, *p*<0.001; **Fig 1B.i**). Combination therapies further lengthened the time to reach 20 mm when compared to olaparib alone (olaparib+alpelisib 115 days versus 75 days, *p*<0.001, **Fig 1B.i**; olaparib+Poly(I:C) 120 days versus 78 days, *p*<0.001, **Fig 1B.ii** and **S1 Fig**). This study demonstrated for the first time that Poly(I:C) enhanced PARP inhibition, with an effect size similar to alpelisib in olaparib-sensitive tumors.

**Fig 1.**
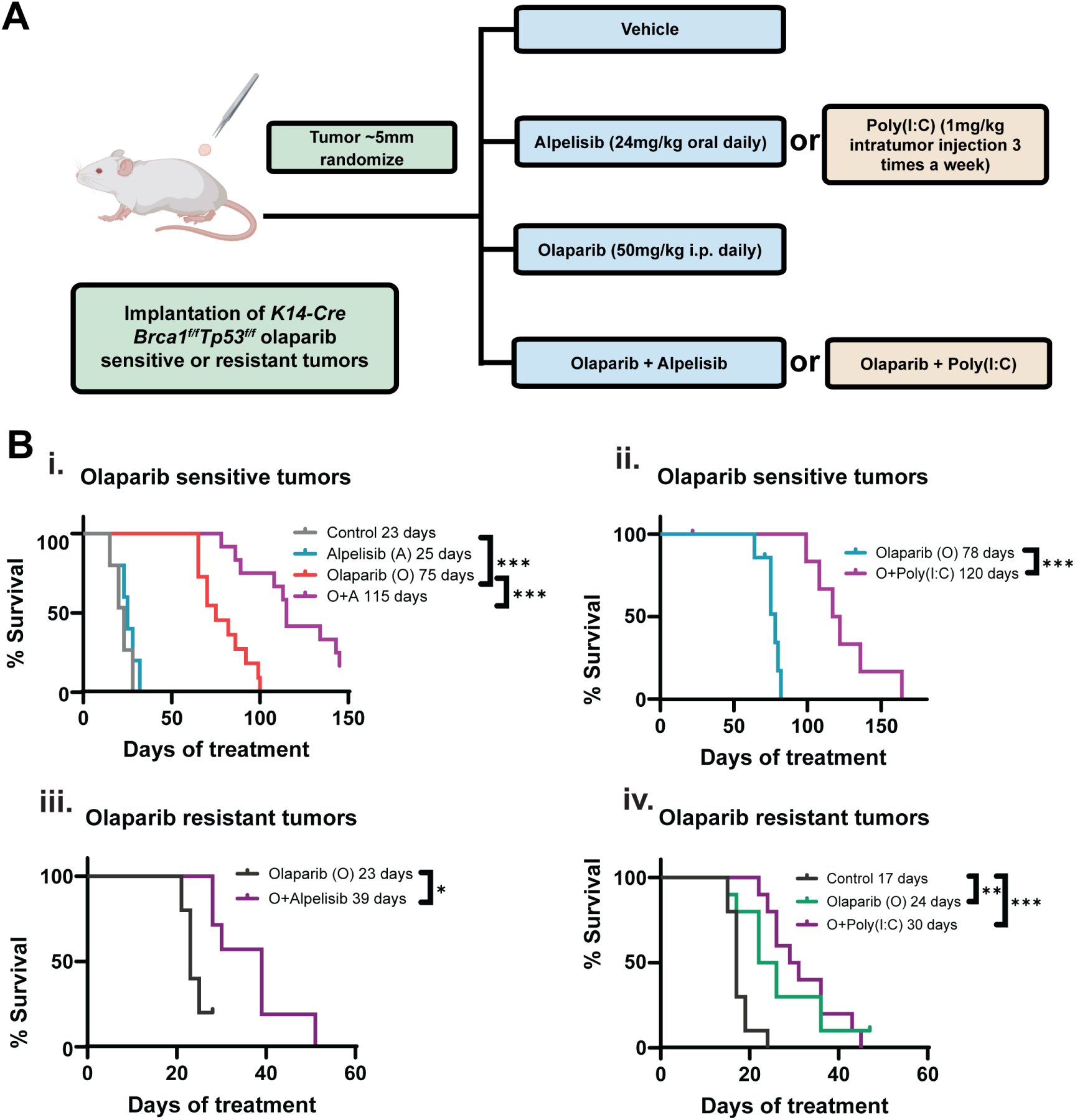
Evaluating combination therapies in a murine *BRCA1*-related TNBC model. **(A.)** The schematic illustrates the *in vivo* study design. Syngeneic olaparib-sensitive or -resistant tumors were implanted into FVB/NJ mice and treated with olaparib alone or in combination with alpelisib or Poly(I:C) once tumors reached a diameter of 5 mm. **(B.)** Kaplan-Meier curves illustrate time taken for tumors to reach study endpoint of 20 mm. The y-axis label “% Survival” denotes the estimated percentage of mice whose tumors had yet to reach the 20 mm endpoint. **i.** Olaparib-sensitive tumors were treated with vehicle (n = 5), olaparib (n = 11), alpelisib (n = 5), or the combination of both (n = 12). **ii.** Olaparib-sensitive tumors were treated with olaparib (n = 7) or a combination of olaparib and Poly(I:C) (n = 7). Olaparib-resistant tumors were treated with **iii.** olaparib alone (n = 7) or in combination with alpelisib (n = 5), and **iv.** vehicle, olaparib alone, or a combination of olaparib and Poly(I:C) (n = 10 per group). Here, n represents the number of mice in each treatment group. Statistical significance is indicated as follows: \**p*<0.05, \*\**p*<0.01, \*\*\**p*<0.001. Created with BioRender.com (panel A schematic only).

We derived olaparib-resistant tumors from initially sensitive tumors grown orthotopically under constant olaparib treatment.[29] Median time to endpoint was shorter in mice with resistant tumors compared to those with sensitive tumors (resistant 23-24 days versus sensitive 75-78 days; **Fig 1B**). Olaparib+alpelisib but not olaparib+Poly(I:C) improved the outcome in resistant tumors when compared to olaparib alone (olaparib+alpelisib 39 days versus 23 days, *p* = 0.018, **Fig 1B.iii**; olaparib+Poly(I:C) 30 days versus 24 days, *p* = 0.567, **Fig 1B.iv** and **S1 Fig**). This suggests that adding alpelisib may improve treatment outcomes despite PARP-inhibitor resistance.

### Spatial miRNA analysis

Given that the combination therapies improved treatment outcomes, we investigated the extent to which different treatments altered the spatial miRNA profiles of the tumors. We performed our previously described hydrogel-based, nanoliter-well *in situ* miRNA assay [14,26,30–32] on tumors (*n*=21) harvested 10 days after treatment when the TME remodeling and transcriptional reprogramming are likely to be complete but not yet confounded by extensive necrosis (**Fig 2A**).

**Fig 2.**
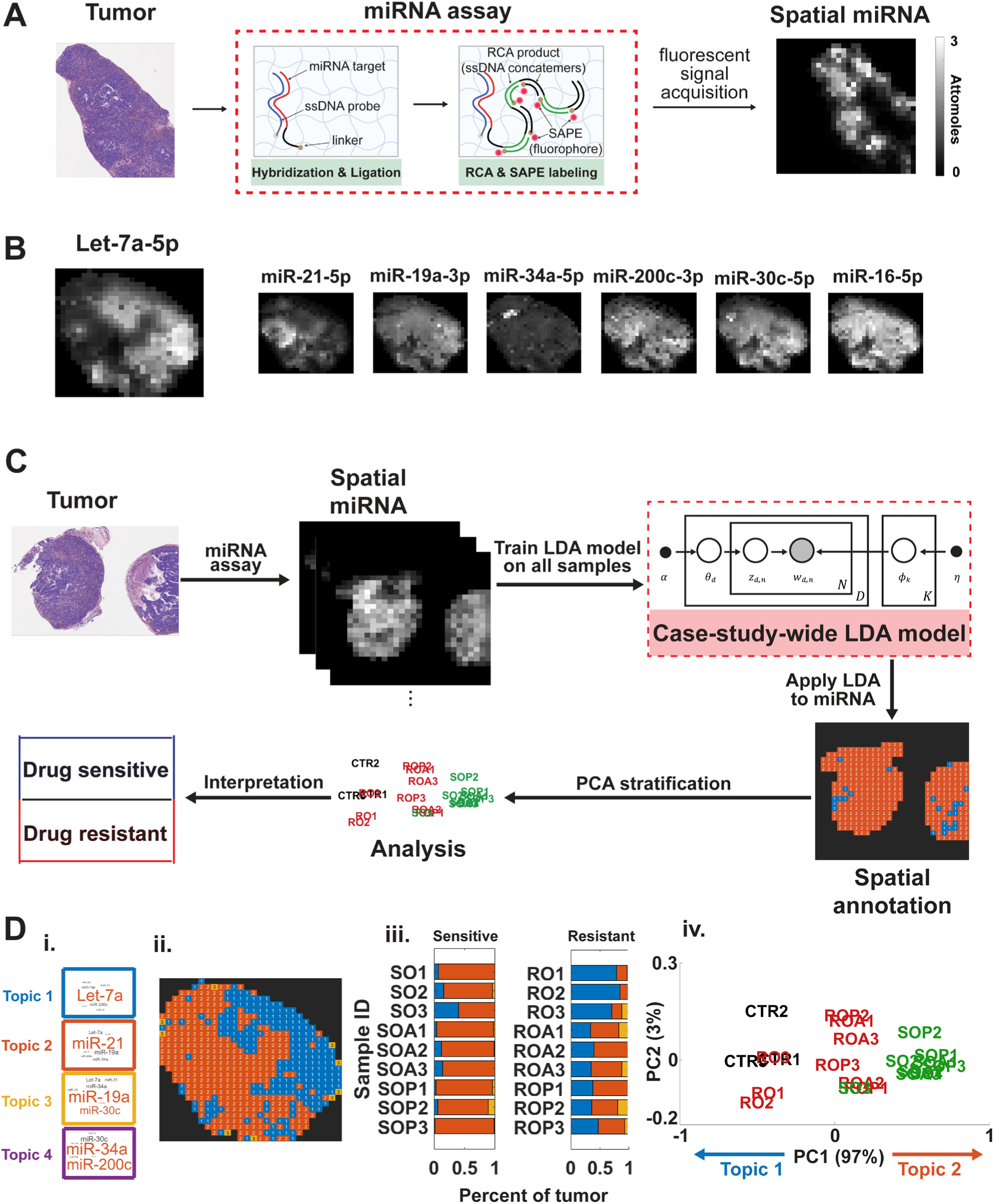
Spatial miRNA analysis overview. **(A.)** Schematic of the miRNA assay used to obtain spatially-resolved miRNA data. **(B.)** Characteristic 7-plex spatial miRNA output. **(C.)** Case-study-wide LDA pipeline for assessing treatment resistance. **(D.) i.** LDA topic results from the case-study-wide LDA model in the form of a word cloud where the font size of each miRNA corresponds to its relative weighting for that topic and the orange font signifies a relatively greater weight within a topic as well. **ii.** The spatial assignment of the LDA topics to a tumor sample; the color of each well corresponds to the color of the topic borders in **i**. **iii.** The percentage of each tissue assigned to each topic. The first letter on the left corresponds to whether a tumor was known *a priori* to be olaparib-resistant (R) or -sensitive (S), followed by treatment group: olaparib-only (O), olaparib+alpelisib (OA), or olaparib+Poly(I:C) (OP); the color of the bars corresponds to the topic border colors. **iv.** PCA of all the tumor samples’ topic percentages. The samples in green correspond to tumors that were known to be sensitive to treatment, while the red denote resistant samples; the black colored text corresponds to control tumor samples. Topic 2 positively contributed towards PC1, while topic 1 contributed negatively.

Our *in situ* miRNA assay simultaneously quantified seven miRNAs commonly studied in breast cancer (**Fig 2B, S2 Fig**).[33–37] MiR-21-5p, let-7a-5p, and miR-19a-3p were highly abundant in these *K14-Cre Brca1^f/f^ Trp53^f/f^*tumors (**S3 Fig**). Cursory univariate analyses determined that both sensitive and resistant tumors treated with olaparib+Poly(I:C) had lower let-7a expression compared to olaparib alone (both adj.*p* = 0.01; **S3 Fig**). The expression of the other six miRNAs did not differ between treatment groups or sensitive/resistant tumors in this bulk analysis (**S3 Fig**). Similarly, a nonspatial tumor-level let-7a/miR-21 ratio did not significantly distinguish sensitive from resistant tumors (Mann-Whitney U test, *p* = 0.546; **S8 Fig**).

Since analysis of individual miRNA expression levels was insufficient to inform tumor phenotype or capture tumor-level miRNA organization, we hypothesized that a spatially informed assessment of the tumors may reveal underlying trends, as evidenced by our prior work.[14] We applied a “naïve” implementation of an LDA-based framework, whereby we trained an LDA model on all the 10-day treated tumors (over 115,248 data points), where each point corresponded to a miRNA amount per nanoliter well in a tumor section. This case-study-wide LDA model learned underlying features in the form of “topics” that were themselves composed of miRNA probabilities across all seven measured miRNAs, which could be visualized as word clouds (**Fig 2D.i**). Of the four learned topics, three contributed materially to the tumor-level compositions, whereas the fourth contributed negligibly (**Fig 2D.iii**; **S5 Fig**). For each well, the fitted LDA model compared the seven measured miRNA amounts with the learned miRNA profiles and assigned the well to the most probable topic (**S4 Fig**). This can be visualized colorimetrically by associating topics with colors that can be assigned spatially, allowing for further examination (**Fig 2D.ii, S4 Fig**). A more holistic analysis can be applied by summarizing the associated topic percentages within all tumor sections (**Fig 2D.iii**) which not only allows easier viewing of topic trends across all samples but can also be used to stratify the data via PCA (**Fig 2D.iv**). The resultant PCA showed that the tumors mostly stratified along the first principal component (PC1, 97% variance explained; **Fig 2D.iv**). Olaparib-sensitive tumors occupied the positive PC1 region, with significantly higher PC1 scores than resistant tumors (median PC1 0.42 vs -0.083; Mann-Whitney U test, p <0.001; Fig 2D.iv).

Topic 2, dominated by miR-21, was found to contribute positively to PC1, while Topic 1, dominated by let-7a, contributed negatively (**Fig 2D.iv**). This means that the miR-21-dominated topic was associated with olaparib sensitivity while the let-7a-dominated topic was associated with resistance. Notably, the sensitivity and resistance phenotypes established by the time-to-endpoint studies **Fig 1** were reflected in the spatial miRNA-based stratification of day-10 tumors (**Fig 2D.ii, S5 Fig**).

Within olaparib-resistant tumors, the case-study-wide LDA model stratified tumors that received olaparib alone and those receiving combination therapies (**Fig 2D.iv**). Combination treatments altered the miRNA profiles of resistant tumors, shifting them toward the profiles of sensitive tumors, specifically by increasing the proportion of the miR-21-dominated topic within the tumors (**Fig 2D.iii**). Within the olaparib-sensitive tumors, there was little stratification between olaparib-only and combination treatment regimens (**Fig 2D.iv**), despite the time-to-endpoint study demonstrating an approximately 1.5-fold increase in median time to endpoint with combination therapies compared with olaparib alone (**Fig 1B**). Together, these data indicate that the miRNA profiles distinguishing olaparib-sensitive from -resistant tumors reflect spatial molecular patterns associated with established sensitivity and resistance phenotypes rather than solely a simple pharmacodynamic response to treatment.

### Refined spatial miRNA analysis of PARP inhibitor-sensitive tumors

Since the case-study-wide spatial LDA analysis (**Fig 2C**) did not provide pharmacodynamic information specific for mono- or combination therapy in olaparib-sensitive tumors as shown in **Fig 1B**, we performed a specialized analysis to improve the stratification of olaparib-sensitive tumors and investigate whether early spatial miRNA profiles could differentiate tumors that received combination therapy versus olaparib-only treatment (**Fig 3A**). Specifically, we examined whether this early stratification could reflect the greater efficacy of the combination regimens observed in **Fig 1**.

**Fig 3.**
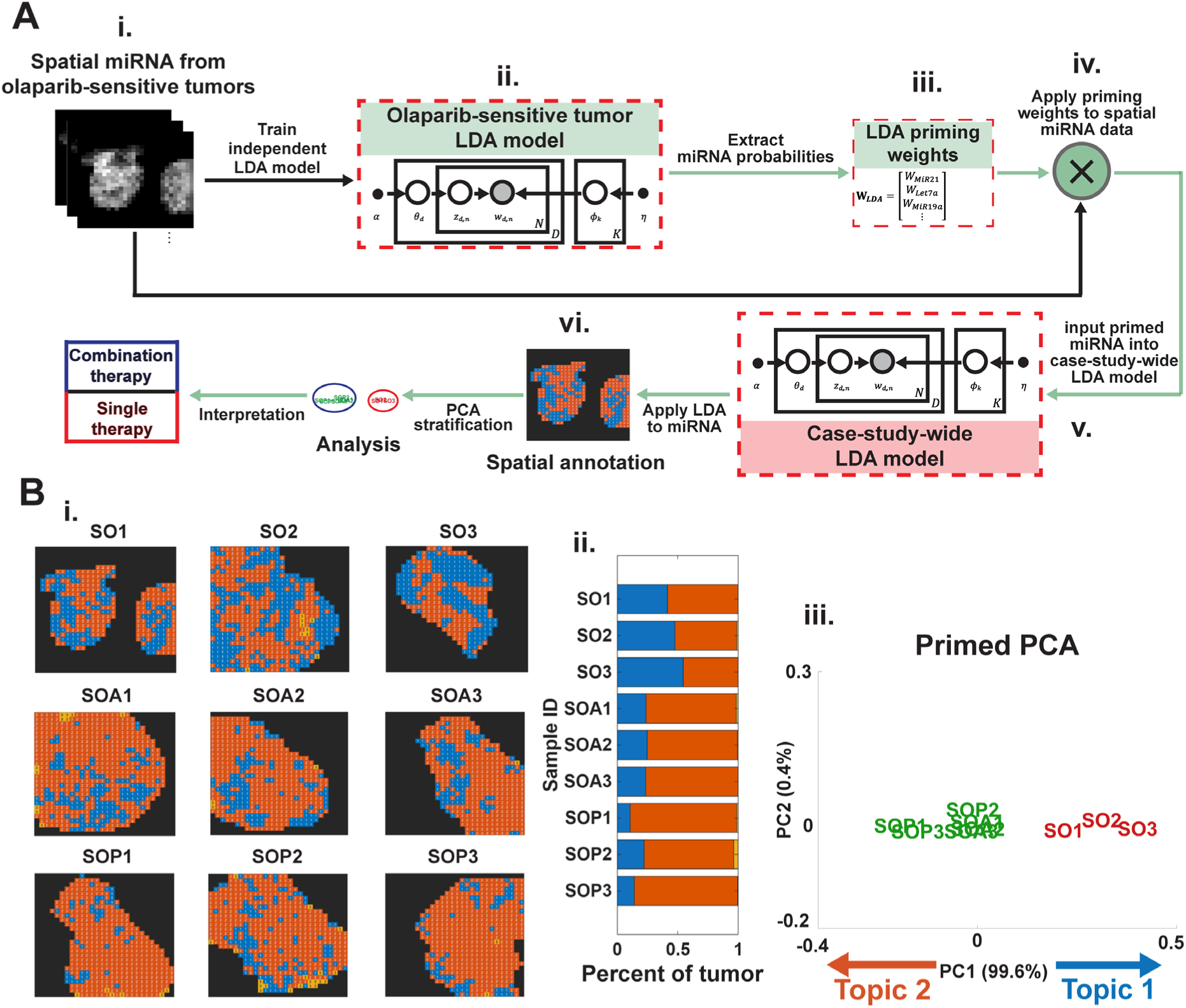
Treatment-sensitive tumor analysis. **(A.)** Schematic of the specialized miRNA analysis of the sensitive-only tumors. **i.** First, the miRNA data of olaparib-sensitive tumors are selected. **ii.** The miRNA data is then used to train an independent LDA model. **iii.** The highest miRNA probabilities from that model were extracted. **iv.** The extracted miRNA probabilities were then used as weights and are multiplied to the input miRNA data, thus “priming” the data. **v.** The case-study-wide LDA model from **Fig 2C** was applied to the primed miRNA data. **vi.** Downstream analysis was performed to stratify the olaparib-sensitive tumors. **(B.) i.** The spatial assignment of the case-study-wide LDA topics to the sensitive tumor samples after being primed by the specialized LDA model miRNA probabilities. The assigned topics were as in **Fig 2D**, where the blue corresponds to the let-7a-dominated topic (topic 1) and orange corresponds to the miR-21-dominated topic (topic 2). **ii.** The percentage of each tissue assigned to each topic. **iii.** PCA of the sensitive tumor samples’ topic percentages. The samples in green correspond to the tumors that received the combination therapies, while the red denote olaparib-only treated samples. Topic 1 was found to contribute positively to PC1, while topic 2 contributed negatively. PC1 scores differed significantly between olaparib-only sensitive tumors (n = 3) and pooled combination-treated sensitive tumors (n = 6; exact two-sided Mann-Whitney U test, *p* = 0.024). A three-group comparison of olaparib-only, olaparib+alpelisib, and olaparib+Poly(I:C) tumors (n = 3 per group) was also significant (Kruskal-Wallis test, *p* = 0.039).

The core of this specialized analysis lies in isolating the sensitive tumor samples and examining how their LDA-derived miRNA probabilities differ from those learned from all the tumors. First, we trained an independent LDA model using only olaparib-sensitive tumors, without incorporating treatment-group labels, to obtain miRNA probabilities (**S2 Table**), which were subsequently used to prime the spatial miRNA input for olaparib-sensitive tumors (**Fig 3A**). The highest miRNA probabilities from the learned topics (**S2 Table**) were used mathematically as weights that were reapplied to the raw miRNA input values, thus biasing the spatial miRNA data with respect to the learned miRNA probabilities (**Fig 3A**). The primed data was input into the original case-study-wide LDA model (**Fig 2C**) without retraining it, and similar downstream visualization techniques and PCA were performed.

The specialized analysis stratified olaparib-sensitive tumors almost exclusively along PC1, with the let-7a-dominated topic 1 now having a greater allocation in tumors treated with olaparib alone, and the miR-21-dominated topic 2 having a greater allocation in tumors that received the combination therapies (**Fig 3B**). The separation along PC1 was statistically significant between olaparib-only sensitive tumors (n = 3) and combination-treated sensitive tumors (n = 6; exact two-sided Mann-Whitney U test, *p* = 0.024). A comparison across the three individual treatment groups also showed a significant difference (Kruskal-Wallis test, *p* = 0.039; Fig 3B.iii). This pattern aligns with the improved longer time to endpoint among sensitive tumors receiving combination therapy (**Fig 1B**) and mirrors the trend seen in olaparib-resistant tumors in which combination treatment similarly shifted the spatial miRNA profile toward the miR-21-dominant topic (**Fig 2D.iii**).

While successful, this stratification still relied on our *a priori* knowledge that the tumors were sensitive and does not establish predictive performance on independent tumors, even though the final topic assignments were allocated by applying the same case-study-wide LDA model as in the naïve analysis to the newly primed miRNA data. What can be gleaned from this specialized analysis, however, is what it reveals about the nature of the spatial miRNA expression that was impacted by the priming and contributed to the successful stratification of the samples by treatment type. In other words, the value of this analysis is not so much that it provides a standalone prognostic classifier as that it reveals which spatial miRNA patterns distinguished olaparib-only from combination therapy within sensitive tumors.

Given that LDA models produce a set of miRNA probabilities that are referred to as “topics,” examining the change between the miRNA probabilities that constitute these topics across both the case-study-wide and the sensitive-only analyses can help explain the change in topic allocation. The highest-probability miRNAs across both LDA models are let-7a and miR-21 (**S2 Table**), which also correspond to the miRNAs that dominate the main topics responsible for stratifying the tumors. Our priming technique utilizes these probabilities as weights applied to the inputs, which results in what effectively amounts to a contrasting effect that is applied spatially across the miRNA data.

In the specialized LDA model, the weights for let-7a and miR-21 are 0.92 and 0.88, respectively (**S2 Table**), while the remaining miRNAs have lower weights. It is those miRNAs with the highest weights that are impacted relatively less than the rest when the LDA model seeks to allocate topics using this newly primed data, such that topics dominated by the reduced miRNAs will have relatively lower values than the topics dominated by heavily weighted miRNAs and can be overtaken by new topic allocations. With respect to our two dominant topics, we notice that, specifically in the olaparib-only treated samples, some of the formerly allocated topic 2 tumor regions, which were miR-21-dominated, are now being assigned to topic 1, which was let-7a-dominated, as can be seen in the spatial annotation maps in **Fig 3B** and **S6 Fig**. Given that the learned let-7a weighting was only ∼4% greater than that of miR-21, the observed reassignment of some regions in the olaparib-only treated samples suggests that the contributions underlying their competing topic assignments were relatively close in magnitude, as opposed to those regions that maintained their original topic assignment. Thus, treatment-type stratification was driven primarily by changes in the spatial topic assignments of these priming-sensitive regions.

### Spatial associations between miRNAs and tumor cellular composition

Although spatial miRNA profiling stratified tumors by olaparib sensitivity and improved our understanding of how relative miRNA abundance related to treatment type, limitations in the current analytical approach remain. The spatial information gleaned from the miRNA data and represented by spatially allocated miRNA-informed topics (**Fig 2D.ii**) was effectively reduced to a summary statistic and further processed via PCA in order to answer a higher-level question: whether tumors could be stratified according to their established sensitivity or resistance to treatment. By stratifying the tumor samples along a PCA with known miRNA topic contributions (**Fig 2D.iv**), we observed within-cohort stratification of tumors according to their established sensitivity and resistance phenotypes. To further characterize these states, we considered that the effectiveness of olaparib depends on the TME [22–24] and that Poly(I:C) is an immune stimulant[25]. We therefore related spatial miRNA-topic patterns to CD45-positive immune-cell infiltration within the TME.

We performed an automated H&E image analysis to classify cells as tumor or stromal. We then quantified tumor-infiltrating immune cells by staining for CD45 on the next serial tumor section cut immediately after the section used for *in situ* miRNA analysis. We extended our case-study-wide LDA analysis pipeline to include an SSIM assessment (**Fig 4A**)[27]. We used the top two miRNA topics dominated by let-7a and miR-21 and the normalized ratio of the spatial distributions of immune cells to tumor cells as input to generate SSIM maps, whereby the structural similarity of each spatial unit within the tumor is assessed by examining its local neighborhood and then calculating the local SSIM between the miRNA topics and cell ratio maps for that neighborhood (**Fig 4A**).

**Fig 4.**
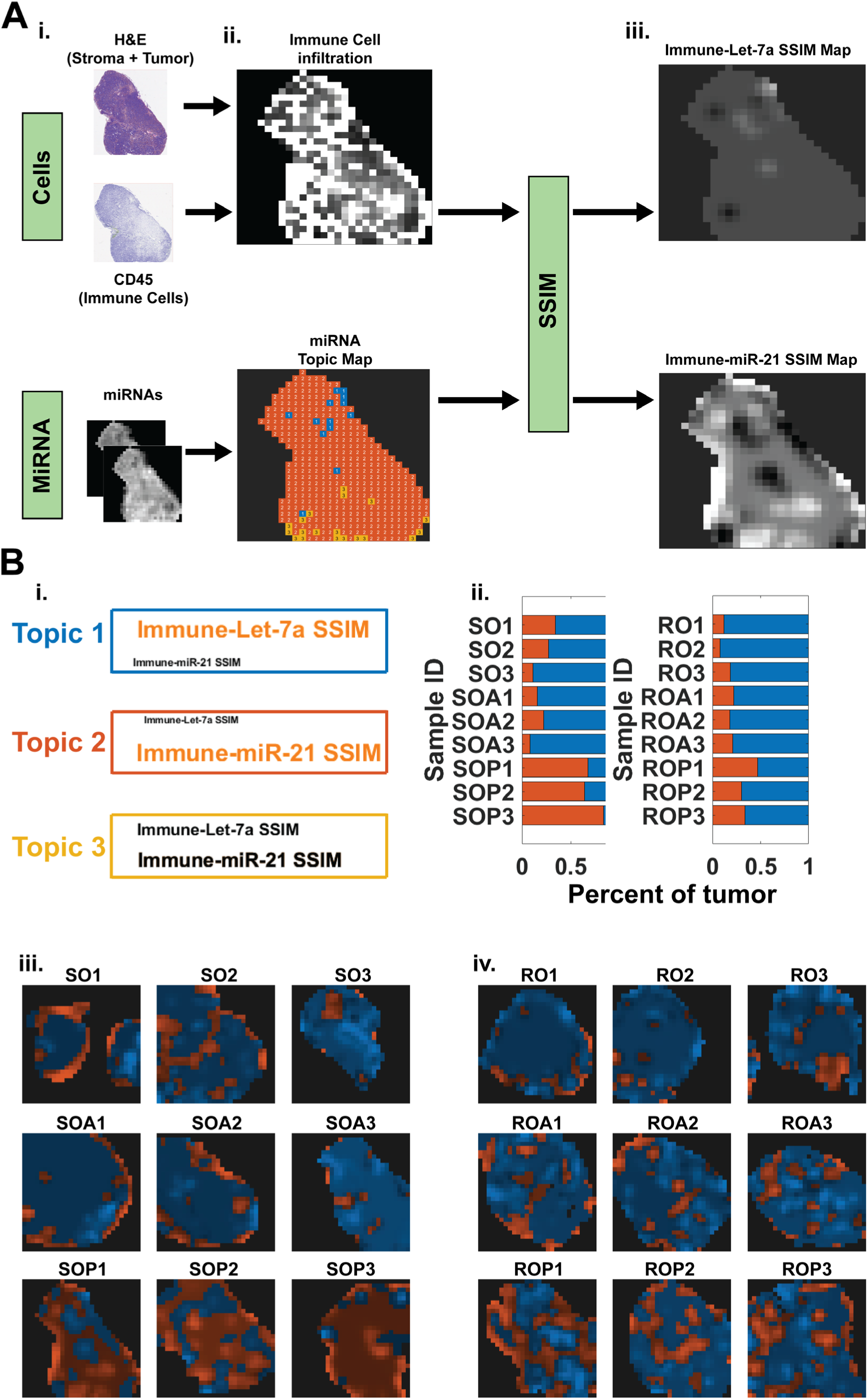
Spatial LDA-SSIM analysis. **(A.)** Schematic of the spatial LDA-SSIM analysis pipeline. **i.** The pre-processed inputs of the analysis pipeline consist of the spatial miRNA data and spatial immune cell counts derived from CD45 immunostaining **ii.** Both the cells and spatial miRNA data were processed prior to input into the SSIM. The miRNAs were processed through the case-study-wide LDA analysis pipeline, and only the top 2 dominant spatial topics were input into SSIM (Fig 2D.ii). Cell counts were processed by calculating the ratio of immune cells to tumor cells, with appropriate normalization applied, to represent spatial immune cell infiltration within each tumor. **iii.** The resulting heatmap representation of the SSIM maps showing the structural similarity between the immune cell infiltration and spatial miRNA topic assignments **(B.) i.** LDA topic results from training an LDA model on the miRNA topic-immune infiltration SSIM maps **ii.** The percentage of each tissue assigned to each SSIM topic. **iii.** The spatial assignment of the SSIM LDA topics to the sensitive samples **iv.** The spatial assignment of the SSIM LDA topics to the resistant samples; the color of each well corresponds to the color of the SSIM LDA topic borders in i. while the intensity of the color corresponds to the intensity of the dominant SSIM maps that inform the topic.

To simplify the multiple per-miRNA-topic SSIM maps into a more interpretable form, we trained a 3-topic LDA model whereby each topic is informed by the miRNA-cell SSIM data. In this model, topic 1 is dominated by the let-7a-immune SSIM, and topic 2 is dominated by the miR-21-immune SSIM (**Fig 4B.i**). This allowed us to determine whether the immune cell infiltration in any given part of the tissue was more similar to the spatial distribution of one miRNA topic as opposed to the other. The 3-topic LDA model was applied to the tumor SSIM maps and the tumors’ topic percentages were calculated (**Fig 4B.ii**). A new variation of the spatial topic map was developed, where we added an intensity scaling to the topic maps to indicate not only which miRNA spatial distributions were more similar to the immune cell infiltration, but also the strength of that similarity via incorporating the per-pixel SSIM scores into the topic map (**Fig 4B.iii, iv**).

Qualitatively, most tumors, regardless of whether they were olaparib-sensitive or - resistant, had a greater spatial similarity between the immune cell infiltration and the let-7a miRNA topic than that of miR-21 (**Fig 4B.ii**). The exception to this observation was found among sensitive tumors that received the combination olaparib+Poly(I:C) treatment, where immune cell infiltration was more spatially similar to the miR-21**-**dominated topic. Given that Poly(I:C) modulates the immune response in tumors [38], increased immune cell activity was expected in tumors receiving olaparib+Poly(I:C). However, CD45 immunostaining and SSIM do not directly measure immune-cell activation. Among olaparib-resistant tumors, mean CD45-positive immune-cell densities were approximately 61.3, 72.3, and 62.7 cells/mm^2^ for olaparib alone, olaparib plus alpelisib, and olaparib plus Poly(I:C), respectively (n = 3 per group). No statistically significant difference was detected among these treatment groups (Kruskal-Wallis test, *p* = 0.646). Further examination of the topic maps revealed a notable pattern in olaparib-sensitive tumors treated with the olaparib + Poly(I:C) combination: miR-21-immune regions appeared non-uniform, with brighter (higher-similarity) signal concentrated near boundaries with let-7a**-**immune regions and dimmer signal toward the center (e.g. SOP3 in **Fig 4B.iii**). Thus, incorporating SSIM into the LDA miRNA analysis pipeline enabled the assessment of local structural similarity between spatial miRNA patterns and the underlying biology. Given the complexity of the spatial analysis, these maps also highlight specific regions that warrant examination in the future.

## Discussion

This study investigated therapeutic strategies combining PARP inhibitors with PI3K inhibitors or the immune-stimulant Poly(I:C) to overcome drug resistance in *BRCA1*-related breast cancers. Both combinations delayed tumor progression, especially in olaparib-sensitive tumors, but that benefit was modest in tumors that had acquired resistance. To evaluate these treatments, we applied our spatial miRNA *in situ* profiling technology to FFPE tumors collected after 10 days of therapy, and developed novel computational frameworks to analyze those early-stage tumors. Our LDA-based analyses demonstrated that early, spatially resolved miRNA profiles distinguished olaparib-sensitive from -resistant tumors, and revealed how combination therapies shifted the spatial miRNA-topic compositions of resistant tumors toward those observed in sensitive tumors. Two spatial miRNA topics, dominated by miR-21 and let-7a, stratified the tumors as olaparib-sensitive or -resistant. By integrating SSIM into the analysis pipeline, we bridged the gap between high-dimensional molecular signatures and the physical tumor cellular composition, enabling examination of local structural similarity between spatial miRNA patterns and immune-cell infiltration. Our miRNA-based spatial analysis shows potential as a useful tool for treatment-response stratification and biomarker discovery, and could ultimately provide a clinical window to adjust therapeutic strategies before tumor progression occurs.

While both PI3K inhibition and Poly(I:C) enhanced olaparib efficacy in sensitive tumors *in vivo*, only the addition of PI3K inhibition significantly prolonged time to endpoint in olaparib-resistant tumors. Our SSIM analysis of tumors after 10 days of treatment showed that, in resistant tumors, there was little difference in the spatial similarity between immune infiltration and the let-7a miRNA topic when comparing olaparib+Poly(I:C) with olaparib alone. The improved treatment outcome with PI3K inhibition suggests that targeting this pathway can provide therapeutic benefit despite acquired resistance, although the underlying mechanisms remain to be established. These findings highlight that spatial miRNA profiling can potentially identify tumors that are sensitive to PARP inhibitors and may benefit from additional immunomodulatory therapies, whereas tumors likely resistant to PARP inhibitors may benefit from the addition of other targeted agents, such as PI3K inhibitors.

We developed a case-study-wide LDA analysis framework that distinguished olaparib-sensitive and -resistant tumors. Among resistant tumors, our initial analysis was able to further distinguish olaparib-only treated tumors and those that had received the combination therapies. The lack of treatment-group separation among olaparib-sensitive tumors prompted us to implement a priming-based, specialized analysis to successfully resolve therapy-associated spatial miRNA profiles. This specialized approach highlighted how targeted priming of spatial miRNA data can reveal subtle stratification patterns that were missed by the case-study-wide LDA analysis.

To understand how priming influenced topic composition, we compared the miRNA probability distributions between the case-study-wide LDA topics and the sensitive-only models. Treatment-specific differences within olaparib-sensitive tumors were encoded in relatively subtle, locally balanced contributions to competing LDA topics, rather than in large global shifts. In both models, the highest-probability miRNAs were let-7a and miR-21 (**S2 Table**), which was consistent with their dominance in the main topics responsible for stratifying the tumors. In the sensitive-only LDA model, the weights for let-7a and miR-21 were 0.92 and 0.88, respectively, while the other miRNAs had lower weights. As a result, those lower-weight miRNAs were reduced more strongly when priming was applied, and topics dominated by the reduced miRNAs lost relative influence during topic assignment compared to topics dominated by the two heavily weighted miRNAs. In olaparib-only treated samples, some of the formerly allocated topic 2 tumor regions (miR-21-dominated) were re-assigned to topic 1 (let-7a-dominated) (**Fig 3B.i vs S5 Fig** and **S6 Fig**). Given that the learned let-7a weighting was only ∼4% greater than that of miR-21, the observed reassignment suggests that the contributions underlying competing topic assignments were relatively close in magnitude in the regions that switched topics, as opposed to regions that maintained their original topic assignment. This explained why priming was able to expose treatment-associated separation within the sensitive tumors even when global shifts appeared modest.

Finally, by integrating SSIM with miRNA topic maps and immune cell infiltration, we not only established a proof-of-concept framework for relating miRNA-defined spatial patterns to cellular architecture, but also found that, whereas most tumors showed greater spatial similarity between immune infiltration and the let-7a-dominated topic, olaparib+Poly(I:C)-treated sensitive tumors instead showed stronger immune-miR-21 spatial similarity. The SSIM-LDA maps showed that immune localization was differentially aligned with distinct miRNA patterns across treatment groups and highlighted specific micro regions where those patterns intersect, providing targets for more detailed pathological and mechanistic follow-up in the future.

The study design should also be considered when interpreting these findings. We focused on a single *BRCA1*-related TNBC mouse model and profiled tumors after 10 days of treatment to capture early spatial miRNA differences before endpoint-associated tissue changes became dominant. Additional treatment stages and BRCA1/2-related tumor models would help test whether the same spatial topics persist beyond this setting. Similarly, validation in independent tumor cohorts would help determine whether the treatment-associated stratification observed following priming can be generalized. The let-7a- and miR-21-dominated topics were associated with olaparib resistance and sensitivity, respectively, but these associations do not by themselves explain the biology underlying those spatial patterns. Although H&E segmentation and CD45 immunostaining characterize the spatial distributions of tumor, stromal, and immune cells, the well-level miRNA measurements do not directly establish the cellular sources underlying the LDA-derived spatial topics. Future studies expanding the seven-miRNA panel and incorporating additional spatial biomarkers and histological analysis of the miRNA-defined regions could help clarify the cellular and molecular features associated with these topics.

Looking forward, these results suggest that spatial miRNA profiling could potentially be leveraged to assess whether a tumor is likely to be responsive to PARP inhibition. Applying our analytical framework to larger preclinical studies and, ultimately, to clinical *BRCA1/2*-mutant breast cancer cohorts would enable the evaluation of the robustness and generalizability of our identified spatial miRNA topics and stratifications, and could facilitate their translation from observed treatment-associated stratification into formal predictive classification models. In parallel, SSIM-based local structural similarity analyses could be extended beyond miRNA-only data to incorporate additional spatially resolved modalities, such as IHC markers and mRNA, and to explore alternative topic models or more complex spatial similarity measures within the same analytical framework, further enriching the biological and translational insight that can be extracted. In the context of PARP inhibitor-based therapy, integrating spatial miRNA readouts with survival outcomes and immune measurements may ultimately help refine patient selection and guide combination therapy choice.

## Methods

### Animal experiments and analysis

All animal experiments were performed in accordance with protocols approved by the Institutional Animal Care and Use Committee at Beth Israel Deaconess Medical Center (protocol number 052-2020-23). Breast tumors generated in *K14-Cre Brca1^f/f^Trp53^f/f^* female mice were dissected into small fragments and transplanted into the mammary fat pad of female FVB/NJ recipient mice from The Jackson Laboratory (RRID:IMSR_JAX:001800) aged ≥6 weeks. The donor mouse line, previously described,[39] was originally provided by the laboratory of Jos Jonkers (Netherlands Cancer Institute, Amsterdam, the Netherlands) and was maintained in-house. Fragments from one olaparib-sensitive donor tumor and one olaparib-resistant donor tumor were implanted into separate recipient mice. Tumors harvested from individual recipient mice were considered biological replicates. Olaparib-resistant tumors were obtained from mice that initially bore olaparib-sensitive tumors and were treated continuously with olaparib until resistance was acquired.[29] For treatment efficacy studies, mice were randomized into treatment groups once tumors reached ∼5 mm in diameter. Treatments continued until tumors reached 20 mm in their longest dimension, which was defined as the experimental endpoint; mice were euthanized at this point. Study arm sample sizes were determined based on prior experience with this tumor model and allowed for possible losses due to unexpected deaths or early euthanasia for tumor ulceration.[18,40] For the 10-day studies, tumors were randomized into treatment groups once they reached 8 mm in size. After 10 days of treatment, tumors were harvested, fixed in formalin, and processed into FFPE sections. A vehicle-only control group was included for the 10-day olaparib-sensitive tumor cohort. A matched 10-day vehicle-only resistant tumor cohort was not included because, when untreated, olaparib-sensitive and olaparib-resistant tumors followed a similar endpoint trajectory, with tumors reaching endpoint within approximately 14-17 days. The time-to-endpoint and 10-day spatial miRNA studies were conducted using separate cohorts of mice. The spatial miRNA study included 21 tumors: three untreated sensitive controls and three biological replicates per treatment group for both sensitive and resistant tumors. Technical replicates of the spatial miRNA assay were not performed in this case study, as assay reproducibility was previously characterized during platform development.[14]

Olaparib was reconstituted in DMSO, diluted in PBS immediately prior to intraperitoneal injection, and administered daily at 50 mg/kg. Alpelisib was administered orally at 24 mg/kg daily. Poly(I:C) was delivered intratumorally at 1 mg/kg, three times per week. Tumor dimensions were measured every 2-3 days using electronic calipers, and tumor volumes were calculated using the ellipsoid formula: Volume=L×W^2^/2, where *L* is the longest dimension and *W* is the shortest dimension. Statistical analyses were performed using GraphPad Prism (RRID:SCR_002798). Growth curves and Kaplan-Meier curves for time to the 20 mm tumor-size endpoint were plotted. Time to endpoint was compared between study arms using log-rank tests.

### H&E and immunohistochemistry (IHC) staining, and automated image analyses

Tumors were harvested and fixed in Rapid-Fixx (Fisher Scientific, Hampton, NH), then submitted to the BIDMC histology core facility (RRID:SCR_009669) for paraffin embedding, sectioning, H&E staining, and performing CD45 IHC using their standard protocol. CD45 IHC was performed using a rat anti-mouse CD45 monoclonal antibody clone 30-F11 at 1:250 dilution (#14-0451-82, ThermoFisher Scientific, Waltham, MA, RRID:AB_467251) and a goat anti-rat IgG horseradish peroxidase polymer secondary antibody at 1:1 dilution (#ab214882, Abcam Inc, Waltham, MA, RRID:AB_3235672). Slides were digitized at 20× using the PhenoImager HT (Akoya Biosciences, Marlborough, MA). QuPath (RRID:SCR_018257) software was used to segment and classify cells as tumor or stromal on H&E images, and quantify CD45+ cells on IHC images.[41,42] For CD45 quantification, the cell-detection classifier was trained using a randomly selected olaparib-treated tumor and subsequently applied to the remaining images. For H&E segmentation, a randomly selected image was used to develop the tissue-segmentation classifier, which was similarly applied to the remaining images to identify tumor and stromal regions. QuPath image analysis was performed without blinding. No slide-specific adjustments were made after classifier setup.

### Spatially resolved miRNA assay

A spatially resolved miRNA assay is applied to tumor samples treated for 10 days to attain the spatial miRNA data. Assay device fabrication and the assay process were performed using methods as described in prior work including use of the same seven-miRNA probe panel.[14] The assay device consists of a nanoliter well array with hydrogel posts embedded within each well. Each well has a 300 x 300 µm footprint and provides a regional measurement of the seven-miRNA panel rather than a single-cell measurement. The hydrogels were functionalized with DNA probes that had complementary sequences to the miRNAs in our panel (**S1 Table**). The relative positioning of the hydrogel posts within each well was designed such that each post in that position would target the same miRNA sequence. The miRNA assay with Rolling Circle Amplification (RCA) was applied to each sample as previously described, and the bound miRNA/DNA complexes were fluorescently labeled which allowed for quantitation.[14] The analytical performance of this assay was previously established,[14] including a detection limit as low as 0.17 zeptomoles and a dynamic range spanning approximately four orders of magnitude.

### Spatial miRNA data pre-processing

The spatial miRNA data acquired from the assay required pre-processing prior to any subsequent analysis (LDA, SSIM, etc.). Utilizing microscope images of the samples during the assay (**S7 Fig**), the portion of the array that was overlapped by tissue, as opposed to the surrounding area outside the sample, was manually outlined and used as a tissue mask so that only miRNA values within the tissue region were retained. The bulk miRNA expression per tumor was compared between the groups using two-way ANOVA using GraphPad Prism. Šídák’s multiple comparisons test was used to compare between two groups, and comparisons that achieved adjusted *p* < 0.05 were reported. For the nonspatial let-7a/miR-21 comparison, the mean expression of each miRNA was calculated across tissue-containing wells for each treated tumor. The ratio of these tumor-level means was log_2_-transformed, and sensitive (n = 9) and resistant (n = 9) tumors were compared using an exact two-sided Mann-Whitney U test (**S8 Fig**).

For LDA model training, the masked spatial miRNA data were processed further in order to ensure optimal performance. First, pixels outside the tissue mask were assigned a value of 0, and an alternate dataset was created in which these masked-out pixels were instead assigned a small positive constant (ε), to provide a safe variant of the data for any logarithmic analyses downstream in the pipeline. Training the LDA model in MATLAB (MathWorks, Natick, MA, RRID:SCR_001622) required restructuring the miRNA data so that the preserved 2D spatial information was linearized by row for all samples and concatenated into a single array for LDA model training. Given this systematic linearization of the data, we were able to transform the results back into the original spatial configuration for visualization and subsequent analysis.

### Case-study-wide spatial LDA implementation

Given the efficacy of the LDA training hyperparameters in prior work [14], an initial 4-topic LDA model was trained on the entirety of the miRNA data derived from all the 10-day treated tumors. The model was trained once using the combined spatial miRNA dataset, without incorporating treatment-response labels, treatment-group labels, or histological information. The learned model was subsequently applied to the individual tumor sections without retraining. Collapsed Gibbs Sampling was used as the inference algorithm, as it has been shown to yield more accurate topic discovery despite taking longer to train (training time is negligible, approximately ∼21 minutes).[43] The model was fit using MATLAB’s fitlda with default options. The fitted Dirichlet priors were symmetric, with parameters of 0.25 per topic and 1/7 per miRNA. The fit met the relative log-likelihood tolerance after five iterations. As opposed to utilizing word frequencies when training the LDA model, we used scaled word counts. The quantified miRNA values per well were on the order of 3-4 attomoles at most and LDA requires integer word counts, so we scaled the values by multiplying every miRNA count by 10^4^ to preserve as much of the data as possible prior to conversion. Once the LDA model was trained, we used the topic word probabilities to visualize the contribution of each miRNA to each topic via word clouds. To assess the stability of the learned topics, the LDA model was refitted five times using different random seeds, and the resulting topic probability distributions were compared with the original model after accounting for topic permutations (**S3 Table**). Applying the trained LDA model on all samples spatially yielded the colored topic maps (**S5 Fig**). The relative abundance of each color-defined topic was then calculated per tumor section to obtain the topic-tumor percentages. Ultimately the LDA output was used to stratify the tumor samples by using the topic-tumor percentages as observations in a case-study-wide PCA, whereby the contribution of the topics (PCA variables) could be noted for each principal component to enable trend interpretation.

### PCA statistical methods

We applied principal component analysis (PCA) directly to the tumor-level LDA topic percentages to visualize how tissue allocation across topics varied among tumors. Each tumor contributed one observation consisting of a vector of topic percentages that summed to 100%. Using the original percentages preserved the interpretation of each input as the fraction of the tumor section assigned to that topic and retained zero allocations where a topic was absent.

For the case-study-wide analysis, scores on the first principal component (PC1) were compared between treated olaparib-sensitive and olaparib-resistant tumors (n = 9 per group) using a two-sided Mann-Whitney U test. For the primed analysis of olaparib-sensitive tumors, PC1 scores were compared between the olaparib-only group (n = 3) and the pooled combination-treatment groups (n = 6) using an exact two-sided Mann-Whitney U test. A Kruskal-Wallis test compared PC1 scores across the three individual treatment groups (n = 3 per group). A *p*-value < 0.05 was considered statistically significant.

### Priming-based specialized LDA analysis for treatment-sensitive tumors

In addition to the case-study-wide LDA workflow, a data priming technique was applied to the sensitive tumors in order to bias the model to stratify them along treatment type, a use that is conceptually consistent with other priming methods in computer vision and other machine learning applications [44–47]. We initially examined all the miRNA probabilities associated with all the LDA topics, then for each miRNA we noted the highest probabilities found across all the topics (**S2 Table**). An auxiliary LDA model was trained using the olaparib-sensitive tumor subset, without incorporating treatment-group labels, to derive the miRNA-specific priming weights. These weights were applied uniformly across all sensitive tumors, and the primed measurements were evaluated using the original fixed case-study-wide LDA model. The miRNA probabilities were then stored in a vector and treated as a form of weights whose values were multiplied to all the spatial miRNA input values for our sensitive tumors. The primed input was then passed through the case-study-wide LDA model and the LDA topics were reassigned and used to stratify olaparib-sensitive samples by treatment type.

### Tumor and immune cell data processing

Tumor and stromal cell count data were obtained from QuPath and by noting the position and orientation of the tumor sections as they were superimposed on our spatial miRNA quantification device while being assayed (**S7 Fig**). Using MATLAB, we quantified the amount of tumor, stroma, and immune cells that occupy the same position as the miRNAs. We defined immune cell infiltration as the normalized ratio of immune cells to tumor cells. First, we defined a tissue mask such that all locations containing immune or tumor cells are included; we then log-transformed the ratio at all locations *(x,y)* while adding a small positive constant ε and where *r(x,y)* represents the new ratio (Equation 1):

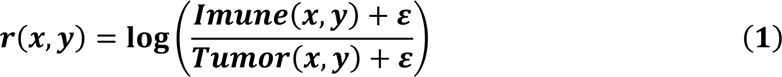

We then remapped the ratios from zero to one via a logistic transform such that values below 0.5 correspond to relatively immune-deficient regions and values above 0.5 correspond to relatively immune-enriched regions (Equation 2):

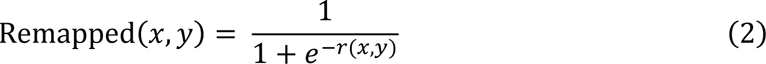

Pixels outside the tissue mask were excluded from downstream normalization. Next, we re-normalized the ratio on a per tumor basis such that the maximum ratio was set to one and the lowest to zero. Lastly, we clipped the values at the median (50^th^ percentile) within the tissue mask and renormalized the clipped values to [0,1]. The reasoning behind the renormalization is to make the final ratio compatible for use with SSIM. Since our second data type (miRNA topic maps) being input to the SSIM pipeline are effectively binarized masks and the spatial similarity between these masks and a ratio where most values are close to zero (given the large difference in immune cell vs tumor cell count), we needed to highlight the significance of the immune cell infiltration via remapping and signal normalization.

### SSIM map formulation

SSIM maps were created in order to encapsulate the structural similarity between our spatial immune cell infiltration ratios and the case-study-wide LDA-derived miRNA topic maps. In essence, SSIM compares two images within a local neighborhood around each pixel by evaluating agreement in local mean intensity, local contrast (variance), and local structure (covariance). This yields a so-called structural similarity index that ranges from -1 to 1, where 1 indicates high local structural similarity, 0 indicates no detectable structural correlation, and -1 indicates strong local anti-correlation (i.e., the structures being compared are inverted relative to one another). We used the built-in SSIM function in MATLAB to formulate the maps with a dynamic range of 1 and a Gaussian standard deviation of 1 pixel (Radius = 1), corresponding to a 7 × 7-pixel Gaussian-weighted neighborhood on the 28 × 28 tumor maps.

### LDA-SSIM implementation

An LDA model was trained on the SSIM maps between the immune cell infiltration ratio maps and the top two miRNA topic maps (let-7a- and miR-21-dominated) in the same manner described for the case-study-wide LDA implementation. The 3 LDA topics learned consisted of probabilities of the two SSIM map types (let-7a-immune SSIM and miR-21-immune SSIM) and were applied spatially and statistically summarized into topic percentages within each tumor and represented via color. The spatial topic assignment was done similarly to the miRNA LDA model but with the added complexity of adding a color depth element to the maps such that the level of structural similarity can be gleaned in addition to which SSIM was most similar for any given region in the tumor.

## Acknowledgements

We gratefully acknowledge funding from the National Institutes of Health (NIH) R01CA235740 to PSD and FJS, R35CA232105 to FJS, R01CA226776 to GMW, R21CA267088 to YJH and GMW. GMW is also supported by the Breast Cancer Research Foundation 24-177. This work was supported in part by the Koch Institute Support (core) Grant P30-CA14051 from the National Cancer Institute. We thank the Koch Institute’s Robert A. Swanson (1969) Biotechnology Center for technical support, specifically the microscopy core. This work was in part supported by a graduate fellowship to ONM from the Ludwig Center at MIT’s Koch Institute for Integrative Cancer Research. This work was in part supported by the Takeda pharmaceutical graduate fellowship to ONM.

## Data Availability Statement

All data and code are publicly available and can be accessed via Zenodo repository: https://zenodo.org/records/24476633

## Conflicts of Interest

GMW declares associated, institutional research funding from Mersana, Gilead, Seagen, Celcuity, Totus Medicines, Pfizer, and Genentech but declares no non-financial competing interests. The other authors declare no competing financial or non-financial interests.

## Supplementary Information

**S1 Fig.**
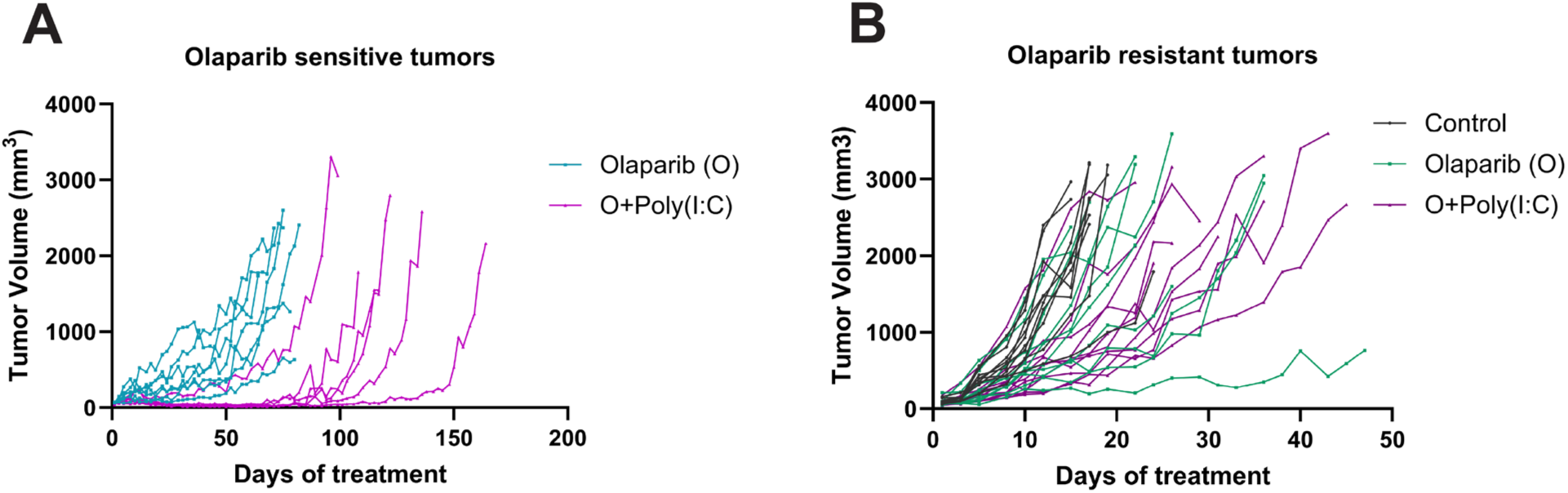
Tumor volume plots. This displays the tumor volume increase over time for the tumors in Fig. 1B. **A.** represents olaparib-sensitive tumors, while **B.** represents the olaparib-resistant tumors.

**S2 Fig.**
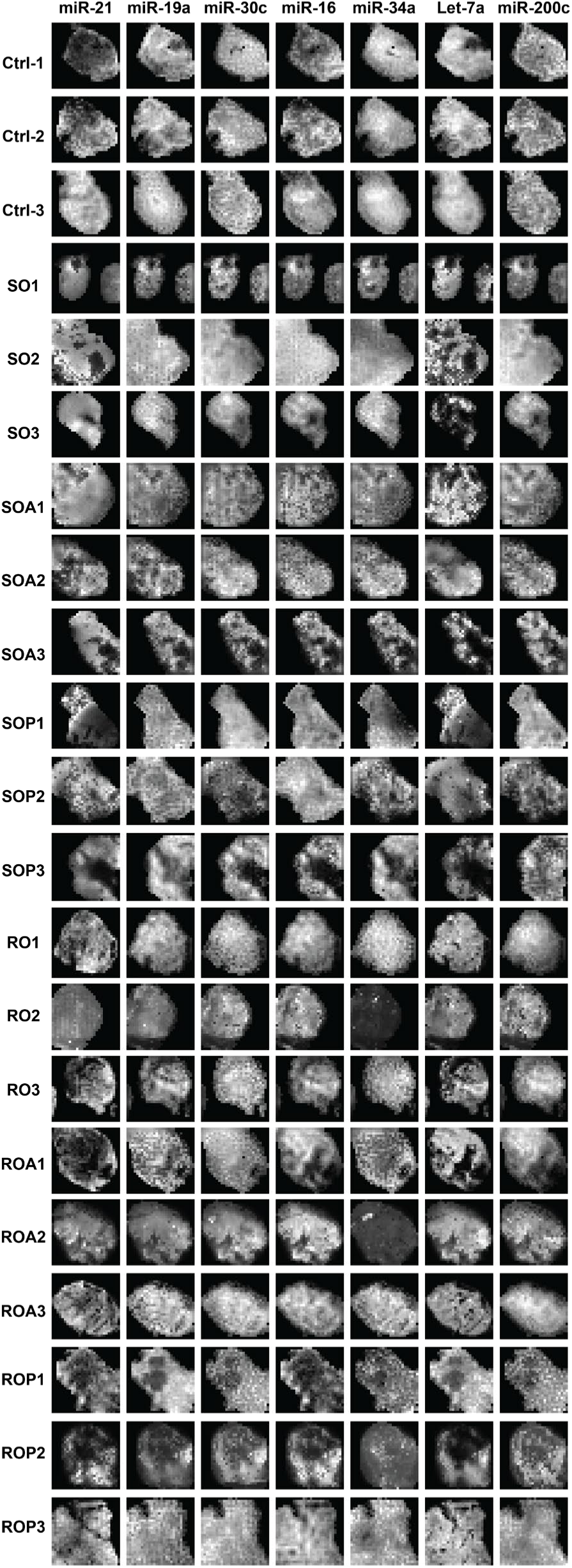
Spatial miRNA dataset. 2D heatmaps of the full data set for all spatial miRNA data for tumors harvested 10 days after treatment.

**S3 Fig.**
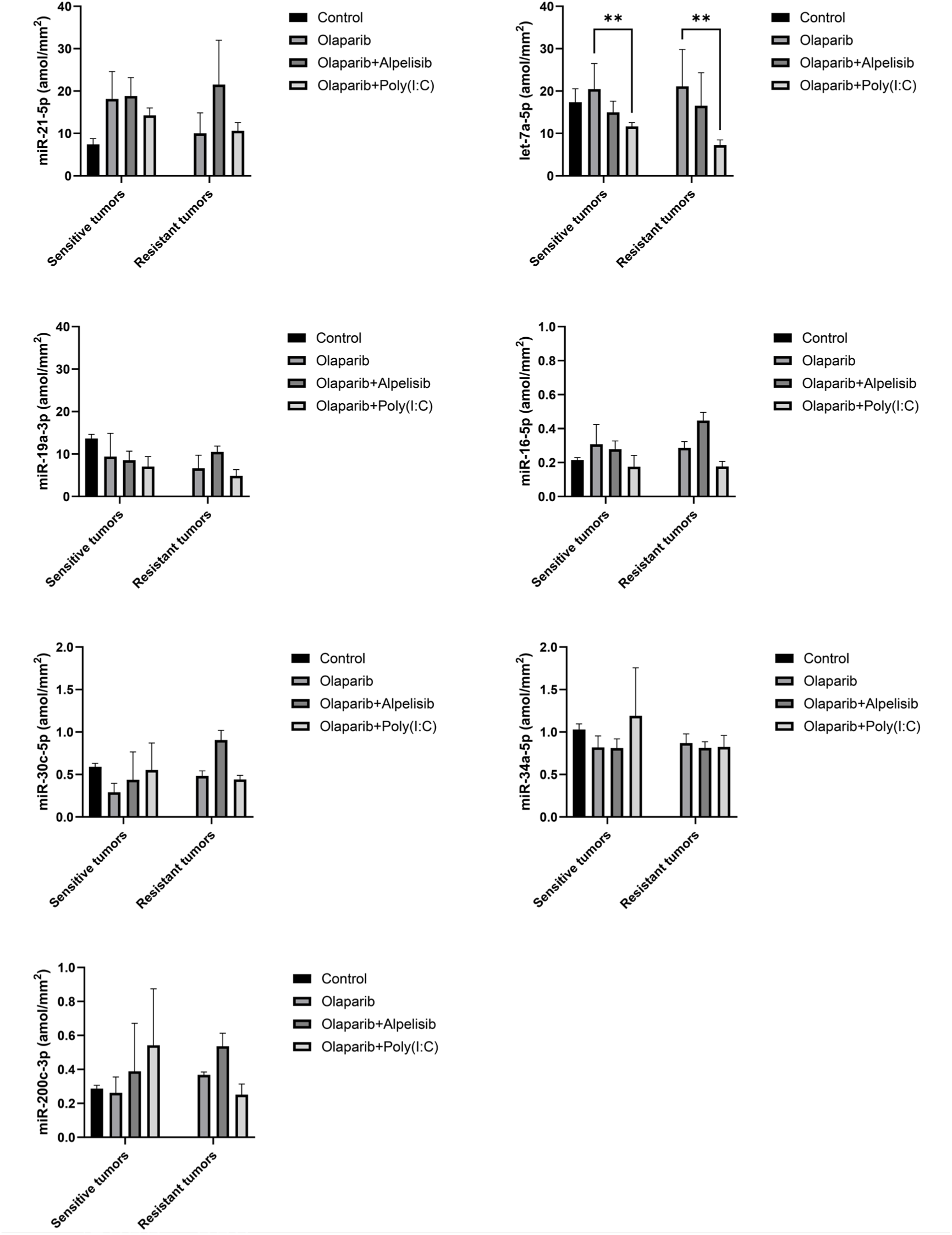
Plots of bulk miRNA expression. The plots for the average miRNA values for all sample types. Each plot represents a single miRNA, the legend indicates the treatment-type administered, and the olaparib-sensitive and -resistant tumors are separated along the x-axis.

**S4 Fig.**
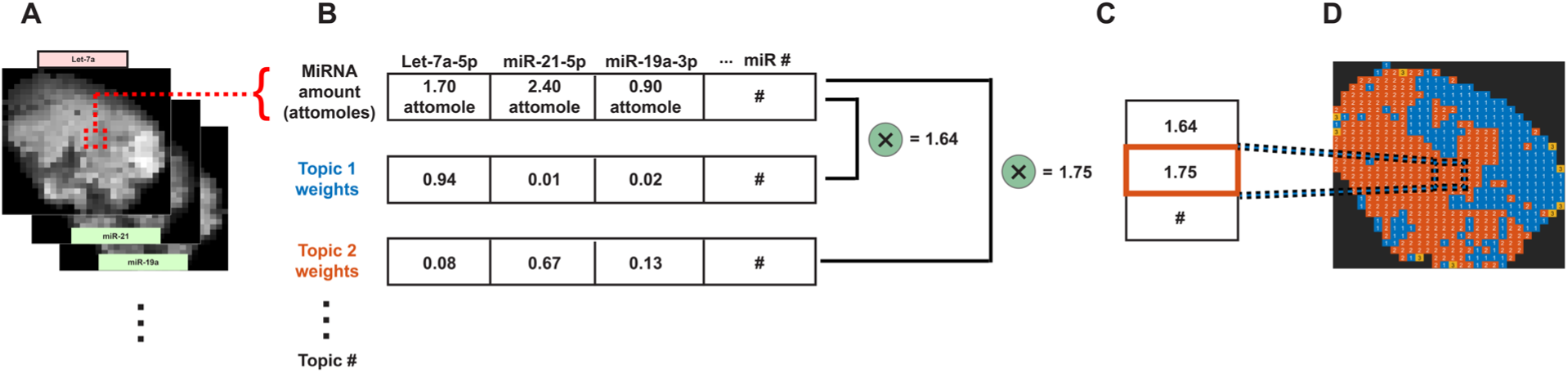
LDA analysis structure. **A.** A tumor’s spatial miRNA data is arranged in 3D slices where for each spatial unit all miRNA values are extracted. **B.** For all the miRNA values, the LDA topic probabilities are multiplied by the miRNA amounts and summed, resulting in a score for each topic. **C.** The topic with the highest score is then spatially allocated to that portion of the tumor and can be depicted colorimetrically as shown in **D**; this process is done for all spatial units for each tumor.

**S5 Fig.**
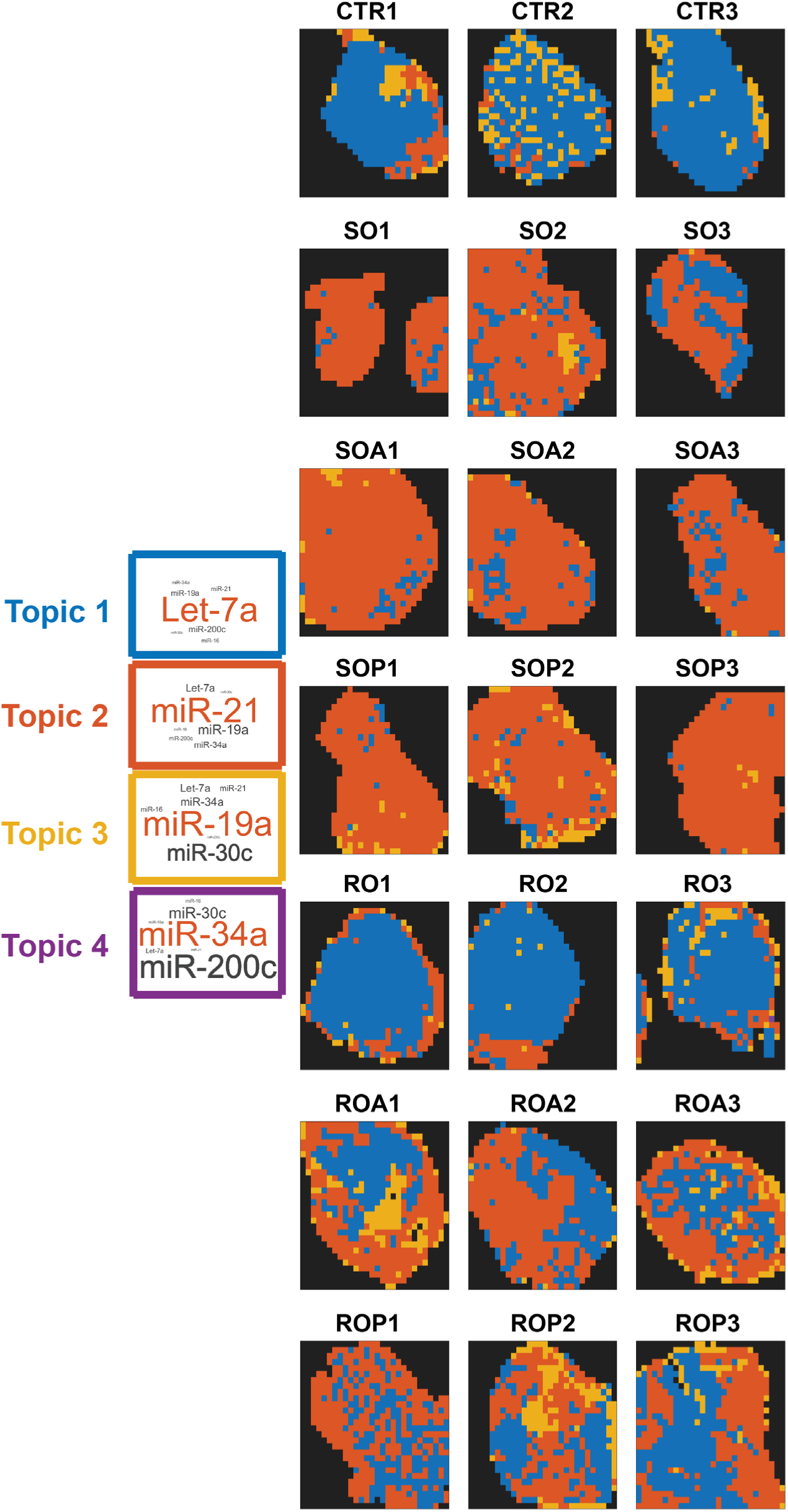
Case-study-wide spatial LDA dataset. The full case-study-wide spatial LDA topic maps for all 10-day tumor samples. The color of the tumors corresponds to the topic bearing the same color.

**S6 Fig.**
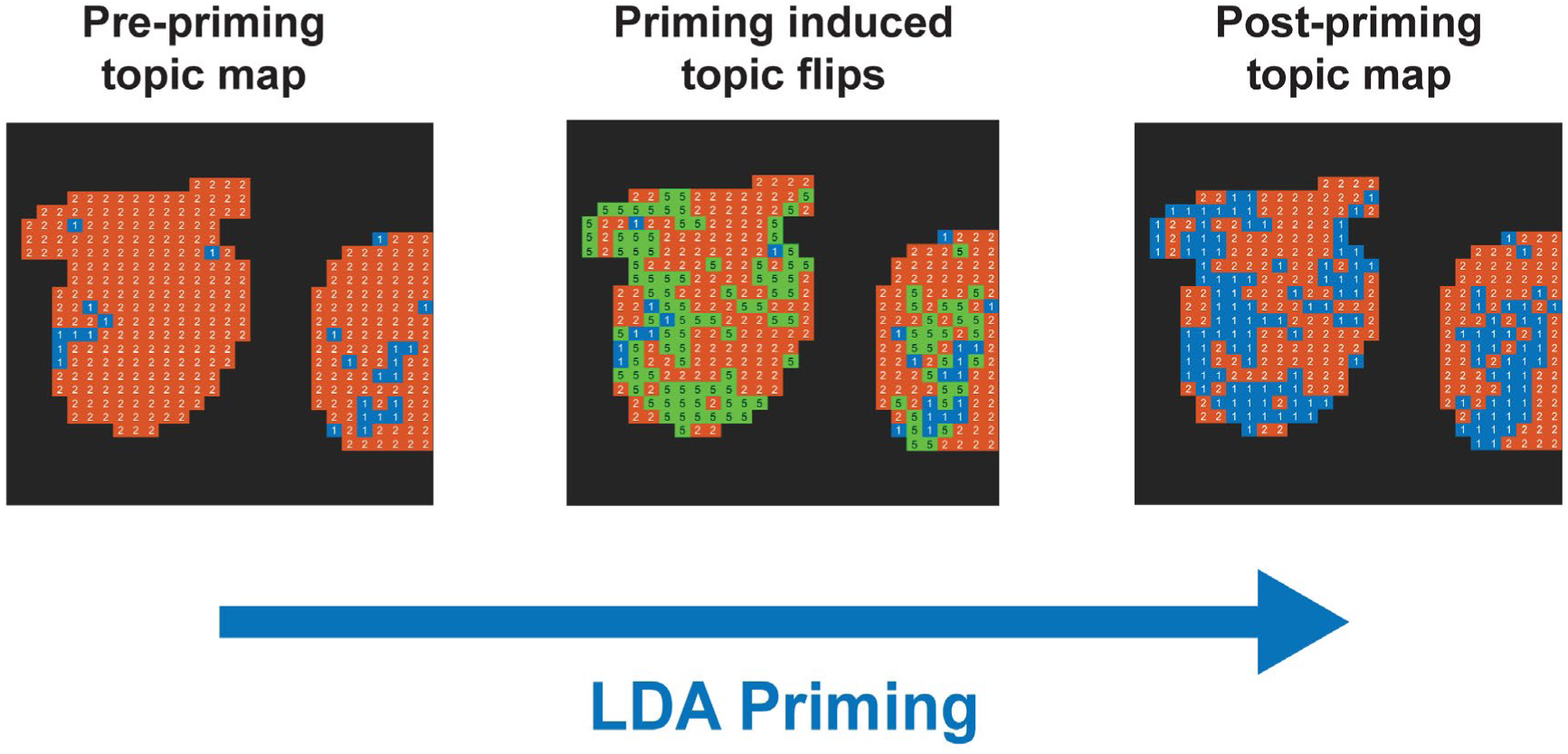
LDA topic changes induced by priming. The application of LDA priming to the miRNA data results in changes (flips) of the determined dominant topic where the pre-biased miRNA amounts of let-7a and miR-21 were close in value. The green represents the locations dominant topic changed as a result of priming.

**S7 Fig.**
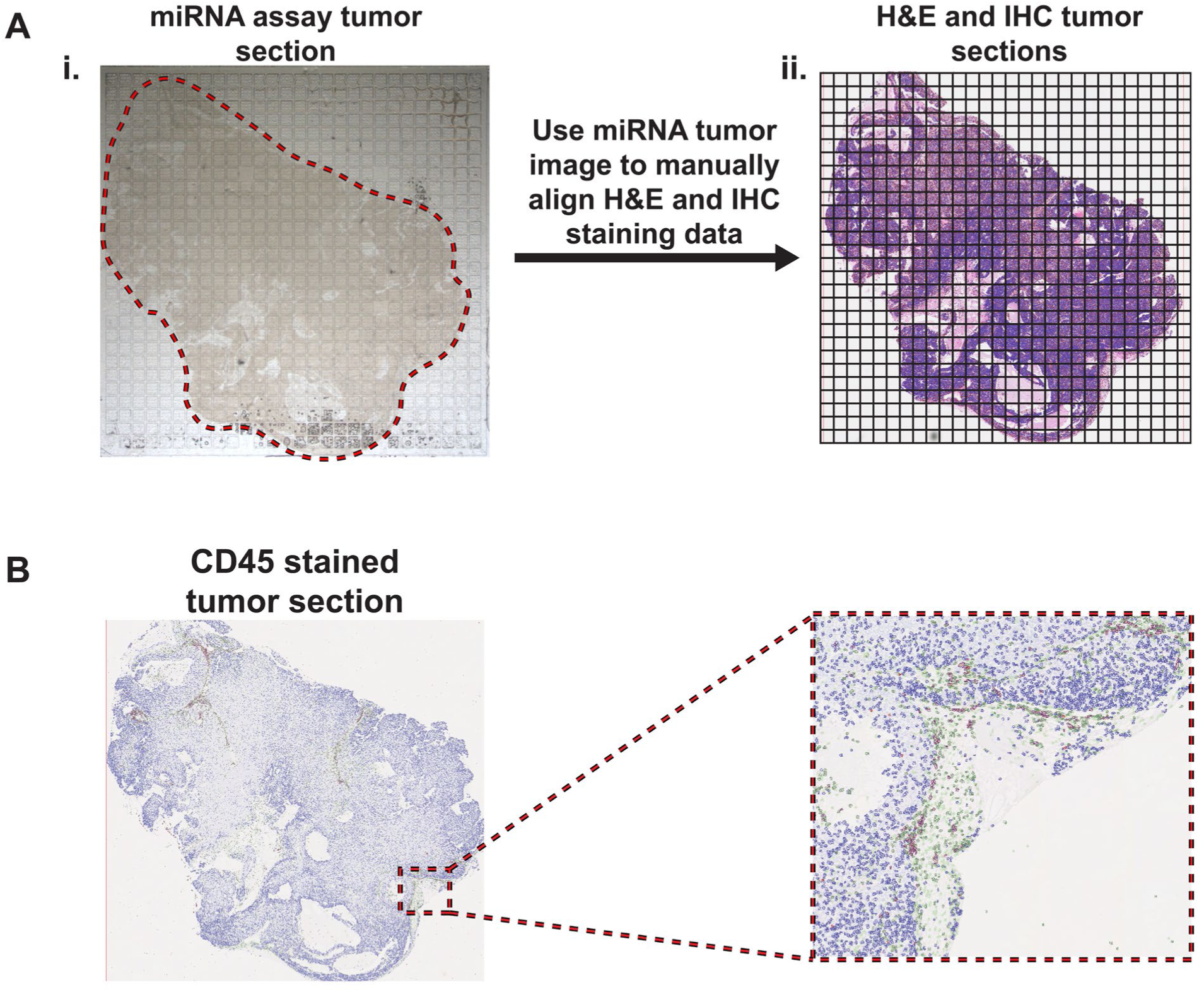
Tumor section alignment and CD45 staining. **A. i.** Tumor sections are superimposed on the miRNA assay device and imaged before running the assay. The captured image is manually outlined to note where the tumor lies within the array. **ii.** A grid with the same dimensions as the assay device is then drawn on images of the adjacent tumor sections that are H&E and stained for CD45 such that they have approximately the same position within the array. This information is used downstream to calculate the number of cells within each well of the assay device via the QuPath derived data. **B.** High quality CD45 stained tumor sections were available for all the 10-day tumors.

**S8 Fig.**
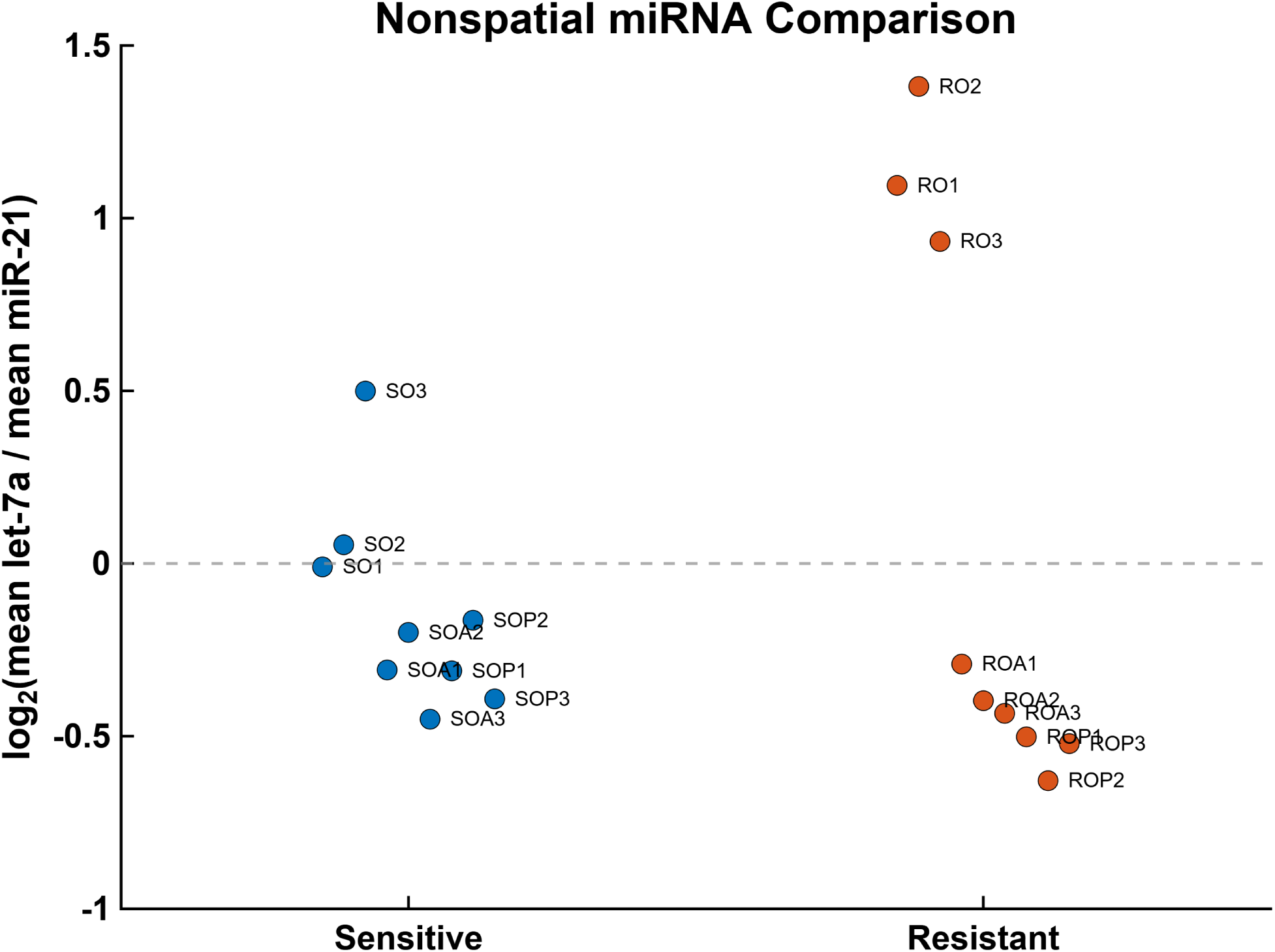
Nonspatial let-7a/miR-21 ratio comparison. Tumor-level ratios of the mean let-7a to miR-21 measurements for olaparib-sensitive (n = 9) and olaparib-resistant (n = 9) tumors. Each point represents an individual tumor, and ratios are displayed on a log2 scale. No significant difference was detected between the two groups (exact two-sided Mann–Whitney U test, p = 0.546).

**S1 Table.**
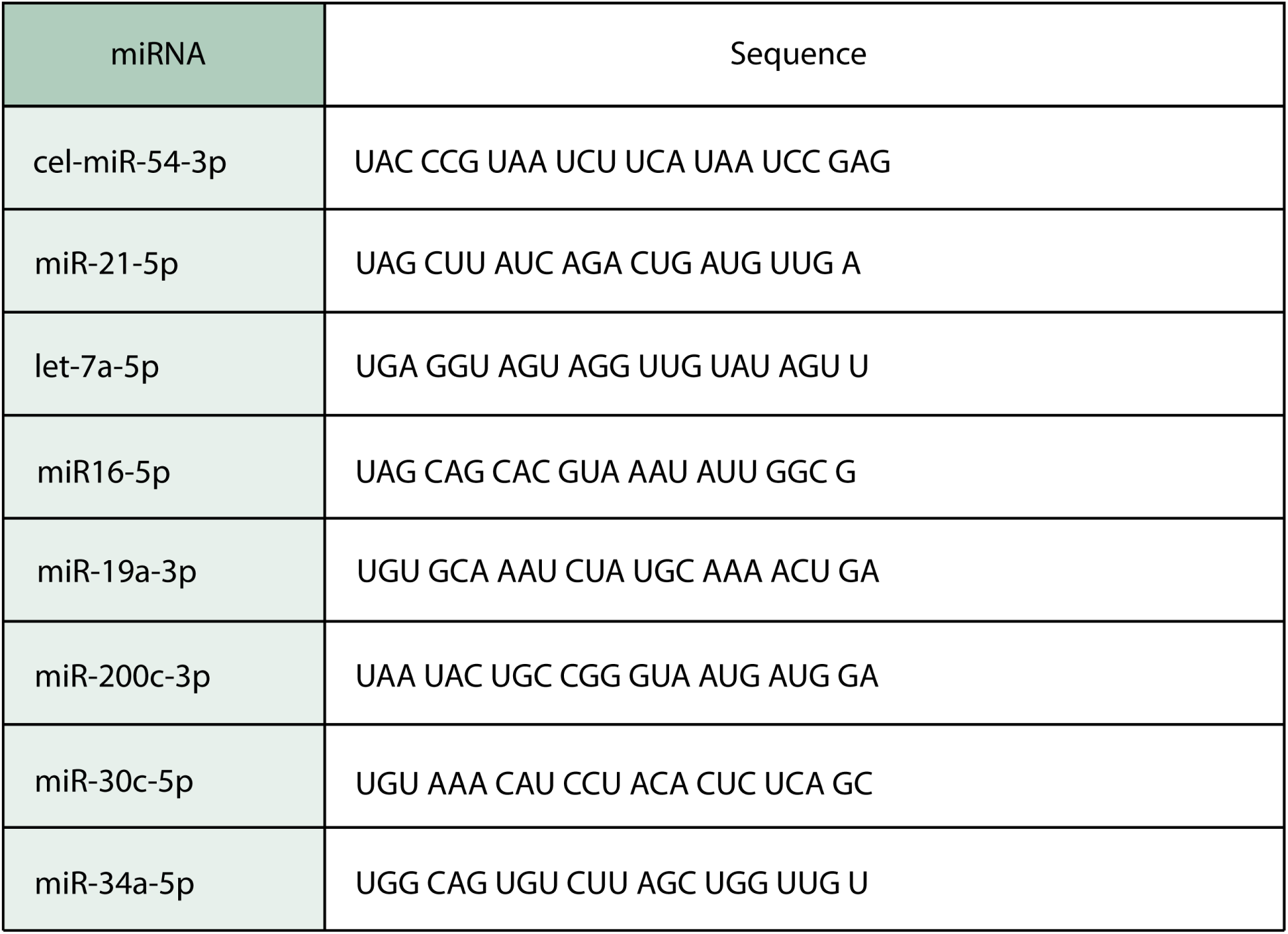
MiRNA sequences used in this work.

**S2 Table.** Top miRNA probabilities across LDA topics. The highest value miRNA probabilities across all topics learned for both the case-study-wide (naïve) and specialized LDA model are noted here. The values for miR-21 and let-7a are the highest across both models. The values in this table obtained from the specialized LDA model are used to prime the miRNA input data for sensitive tumor analysis.

| miRNA | Largest miRNA probability across naïve LDA model topics | Largest miRNA probability across specialized LDA model topics |
|---|---|---|
| miR-21-5p | 0.67 | 0.88 |
| let-7a-5p | 0.94 | 0.92 |
| miR-19a-3p | 0.60 | 0.84 |
| miR16-5p | 0.02 | 0.50 |
| miR-200c-3p | 0.41 | 0.34 |
| miR-30c-5p | 0.26 | 0.04 |
| miR-34a-5p | 0.48 | 0.05 |

**S3 Table.**
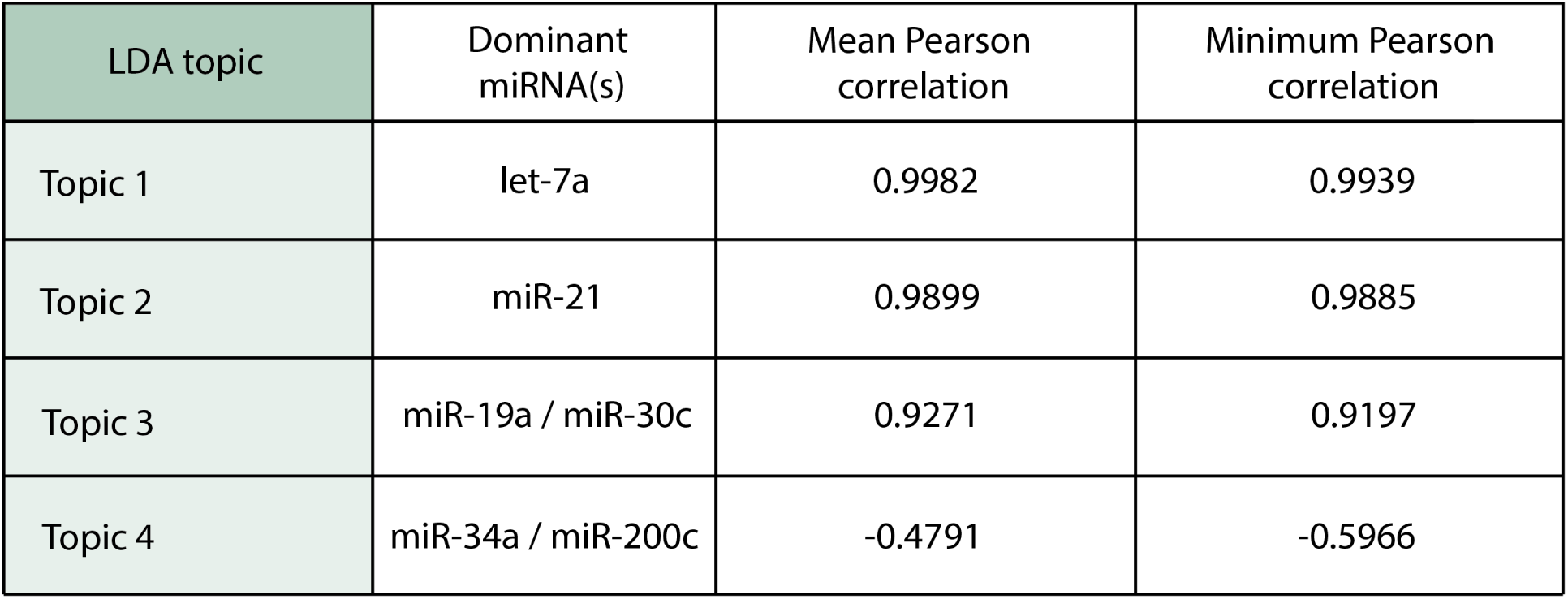
Stability of LDA topic probability distributions across random seeds. The case-study-wide LDA model was refitted five times using different random seeds (1-5). The resulting topics were matched to the original model to account for arbitrary topic-number permutations. Pearson correlations were calculated between the corresponding seven-miRNA topic probability distributions. Mean and minimum correlations across the five runs are reported. Topics 1-3, which contributed materially to the tumor-level topic compositions, were highly reproducible, whereas Topic 4 contributed negligibly and exhibited substantially lower stability.

| LDA topic | Dominant miRNA(s) | Mean Pearson correlation | Minimum Pearson correlation |
|---|---|---|---|
| Topic 1 | let-7a | 0.9982 | 0.9939 |
| Topic 2 | miR-21 | 0.9899 | 0.9885 |
| Topic 3 | miR-19a / miR-30c | 0.9271 | 0.9197 |
| Topic 4 | miR-34a / miR-200c | -0.4791 | -0.5966 |

## Notes

https://zenodo.org/records/18499891

### Summary of Updates

We revised the manuscript in response to peer review, clarified the methods and interpretation of the findings, and added statistical analyses and model stability results to the manuscript and Supplementary Information.

