## Supporting Information for "MicroRNA spatial profiling for assessing drug efficacy in *BRCA1*-related triple-negative breast tumors"

**Contents**

S1 Table. MiRNA sequences in this work

S2 Table. Top miRNA probabilities across LDA topics

S3 Table. Stability of LDA topic probability distributions across random seeds

S1 Fig. Tumor volume plots

S2 Fig. Spatial miRNA dataset

S3 Fig. Plots of bulk miRNA expression

S4 Fig. LDA analysis structure

S5 Fig. Case-study-wide spatial LDA dataset

S6 Fig. LDA topic changes induced by priming

S7 Fig. Tumor section alignment and CD45 staining

S8 Fig. Nonspatial let-7a/miR-21 ratio comparison

**S1 Table. MiRNA sequences used in this work**.

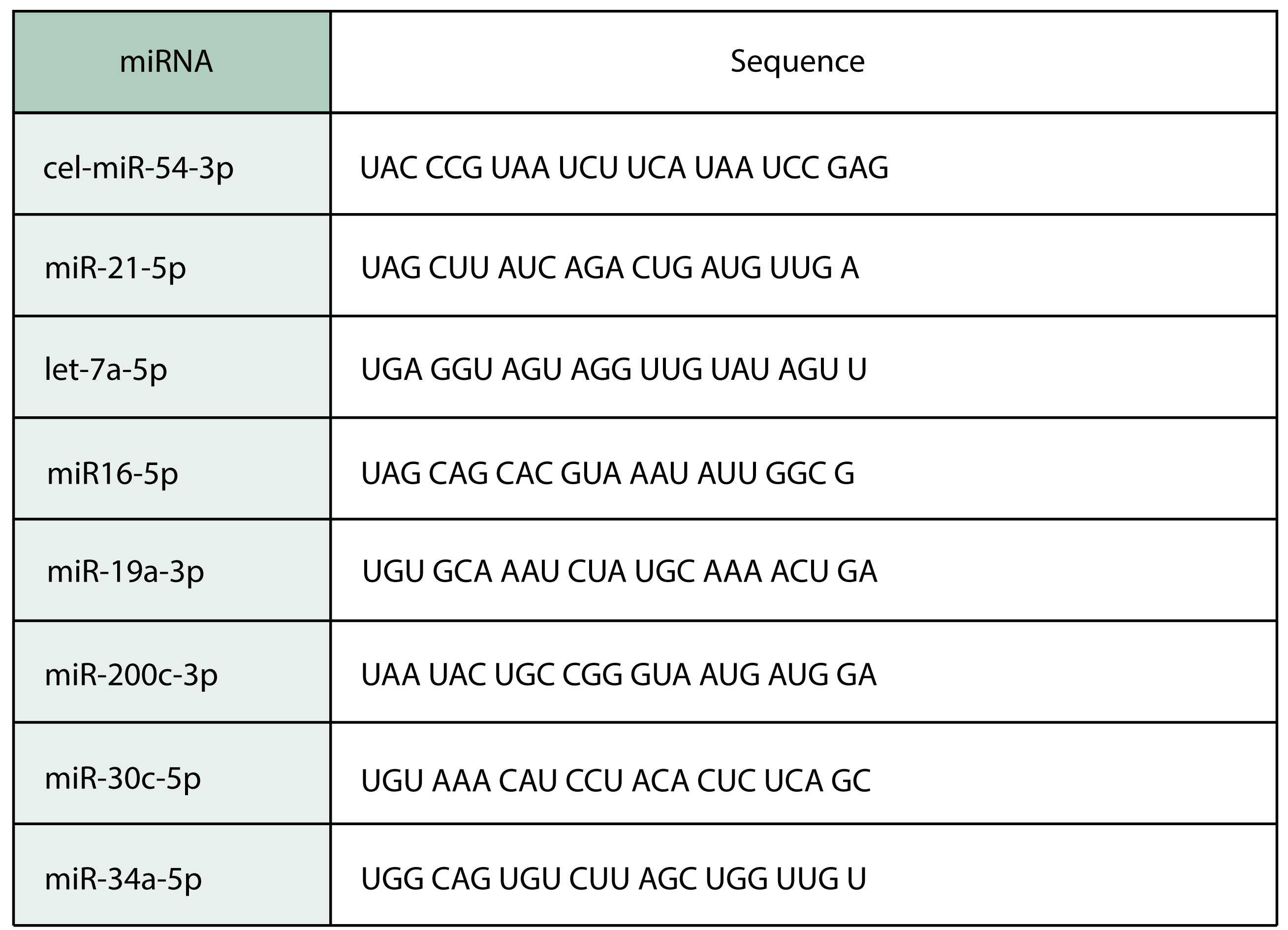

**S2 Table. Top miRNA probabilities across LDA topics**. The highest value miRNA probabilities across all topics learned for both the case-study-wide (naïve) and specialized LDA model are noted here. The values for miR-21 and let-7a are the highest across both models. The values in this table obtained from the specialized LDA model are used to prime the miRNA input data for sensitive tumor analysis.

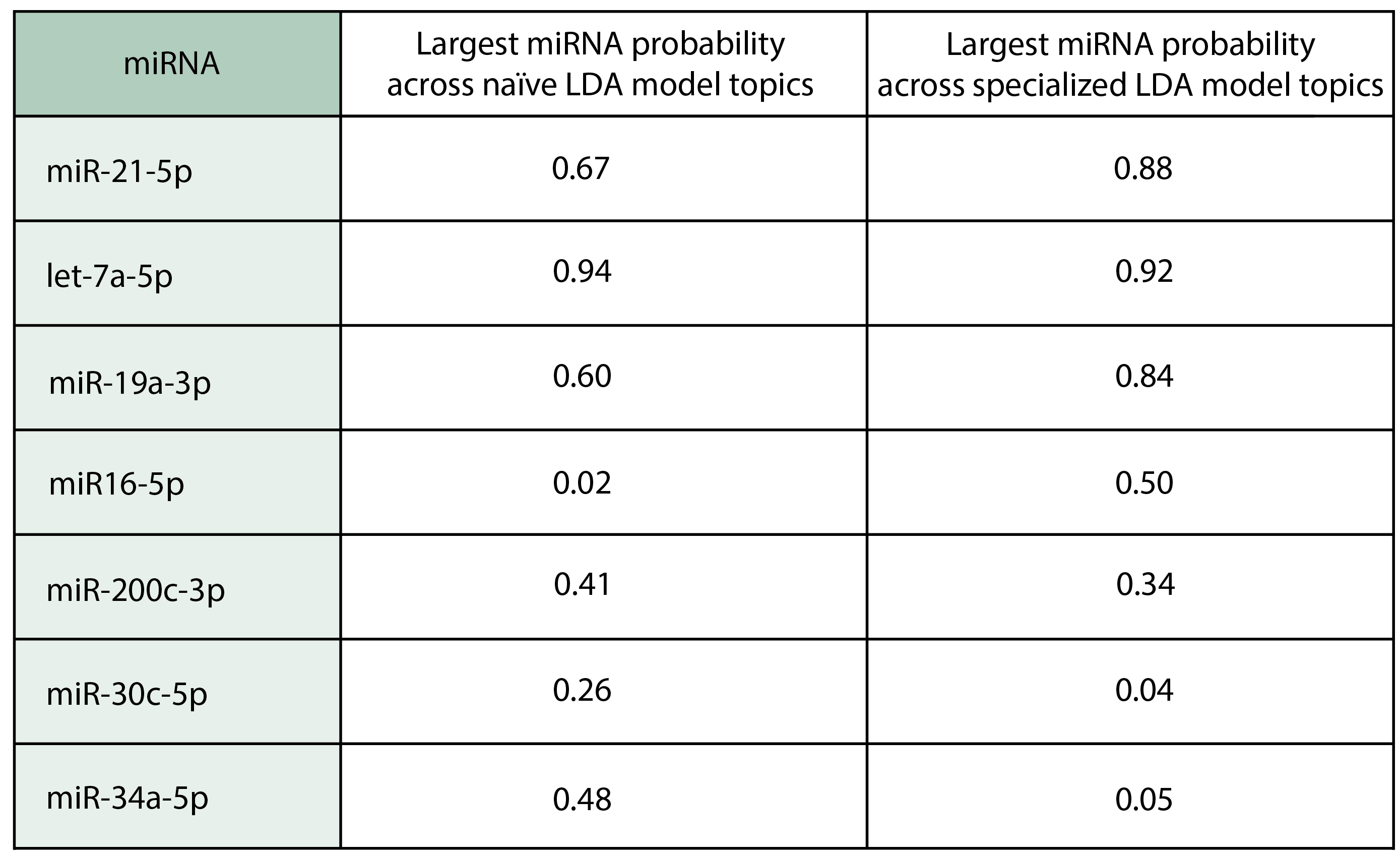

**S3 Table. Stability of LDA topic probability distributions across random seeds**. The case-study-wide LDA model was refitted five times using different random seeds (1-5). The resulting topics were matched to the original model to account for arbitrary topic-number permutations. Pearson correlations were calculated between the corresponding seven-miRNA topic probability distributions. Mean and minimum correlations across the five runs are reported. Topics 1-3, which contributed materially to the tumor-level topic compositions, were highly reproducible, whereas Topic 4 contributed negligibly and exhibited substantially lower stability.

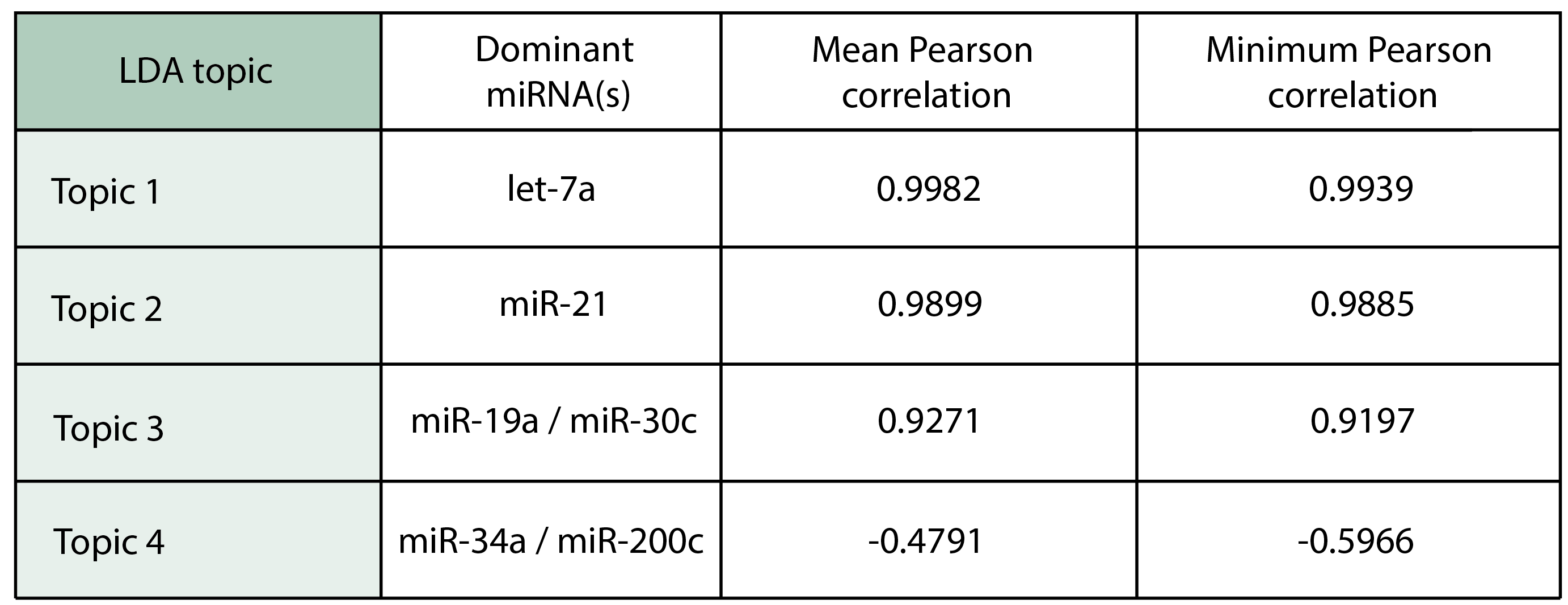

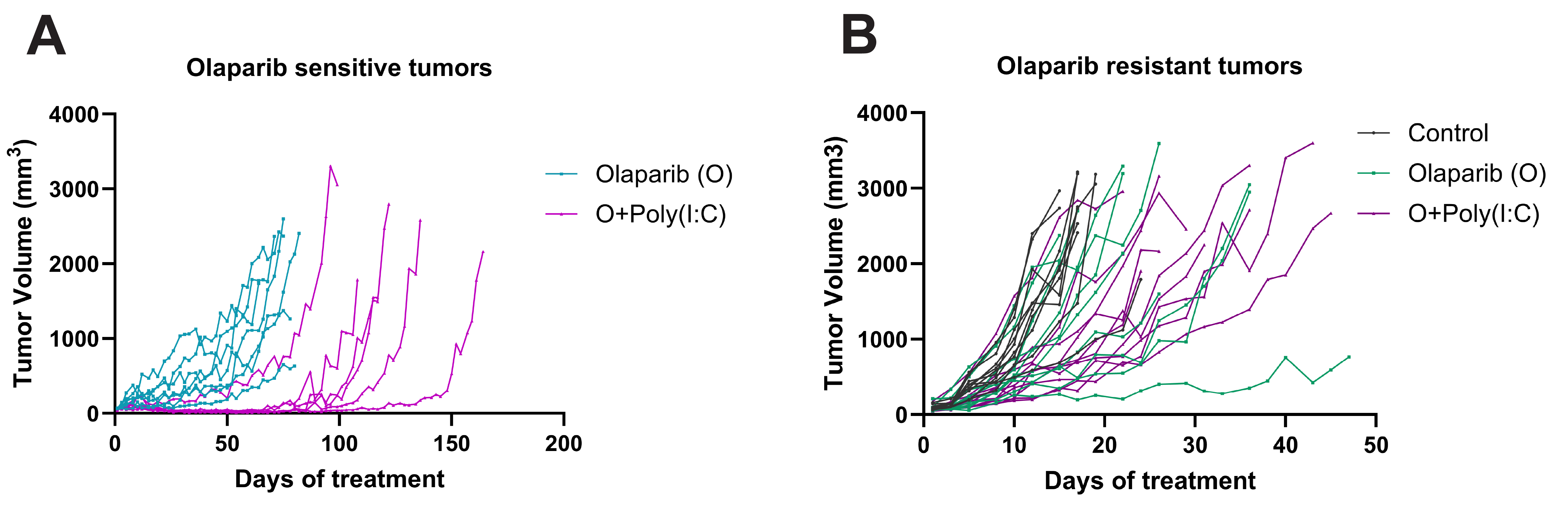

**S1 Fig. Tumor volume plots**. This displays the tumor volume increase over time for the tumors in **Fig. 1B**. **A.** represents olaparib-sensitive tumors, while **B.** represents the olaparib-resistant tumors.

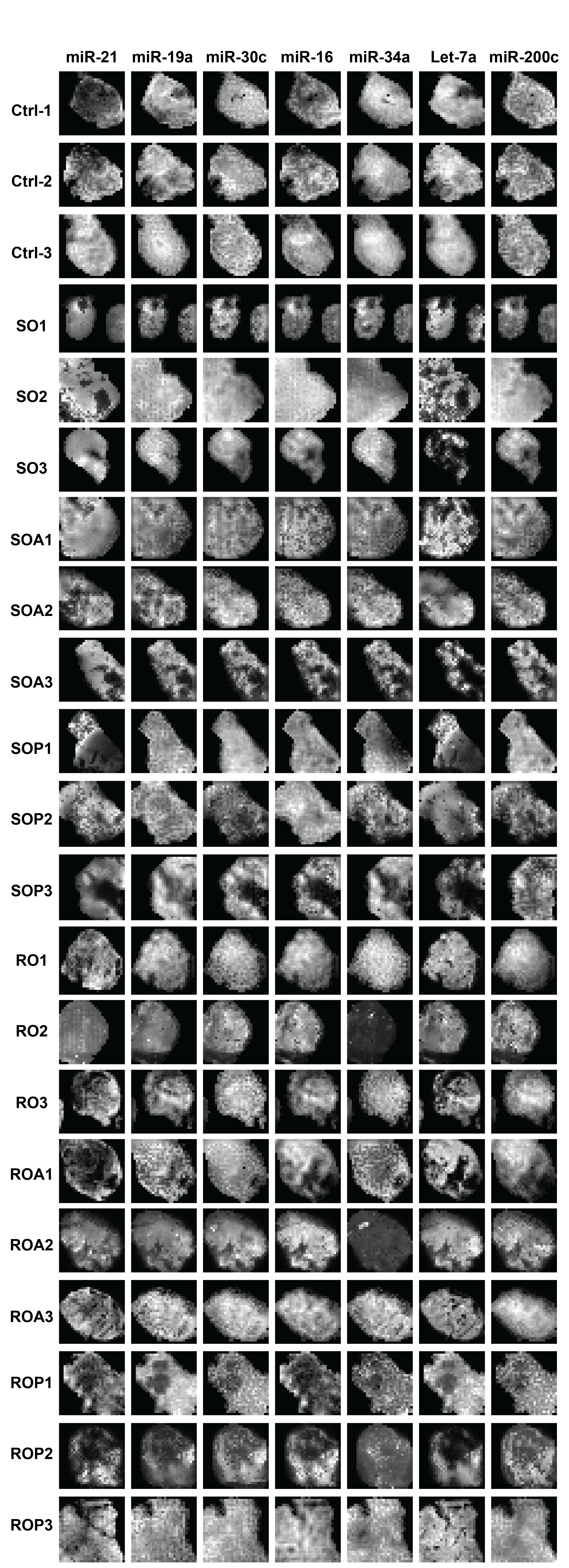

**S2 Fig. Spatial miRNA dataset**. 2D heatmaps of the full data set for all spatial miRNA data for tumors harvested 10 days after treatment.

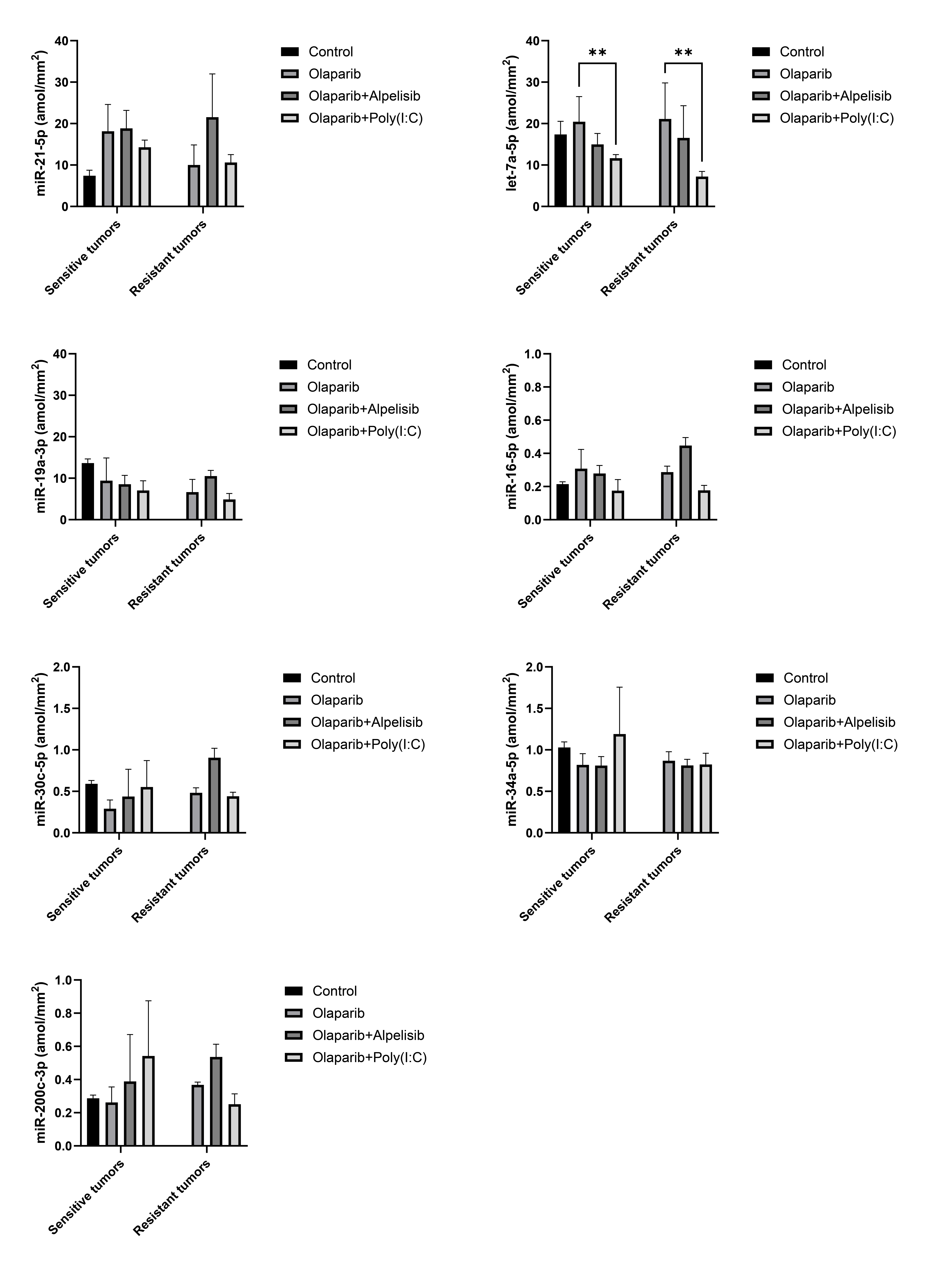

**S3 Fig. Plots of bulk miRNA expression**. The plots for the average miRNA values for all sample types. Each plot represents a single miRNA, the legend indicates the treatment-type administered, and the olaparib-sensitive and -resistant tumors are separated along the x-axis.

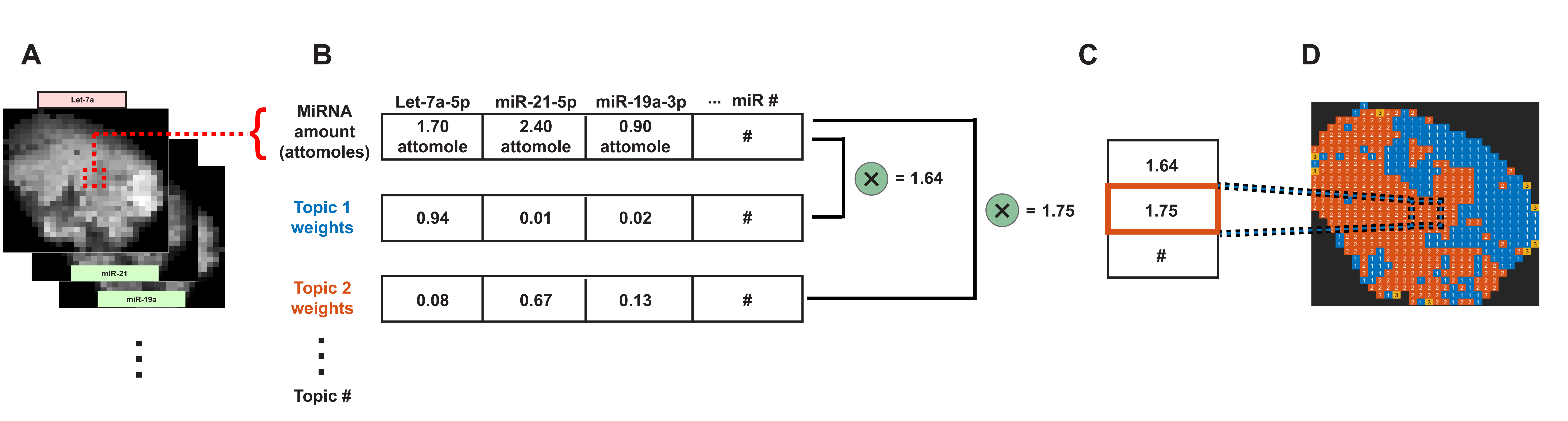

**S4 Fig. LDA analysis structure**. **A.** A tumor’s spatial miRNA data is arranged in 3D slices where for each spatial unit all miRNA values are extracted. **B.** For all the miRNA values, the LDA topic probabilities are multiplied by the miRNA amounts and summed, resulting in a score for each topic. **C.** The topic with the highest score is then spatially allocated to that portion of the tumor and can be depicted colorimetrically as shown in **D**; this process is done for all spatial units for each tumor.

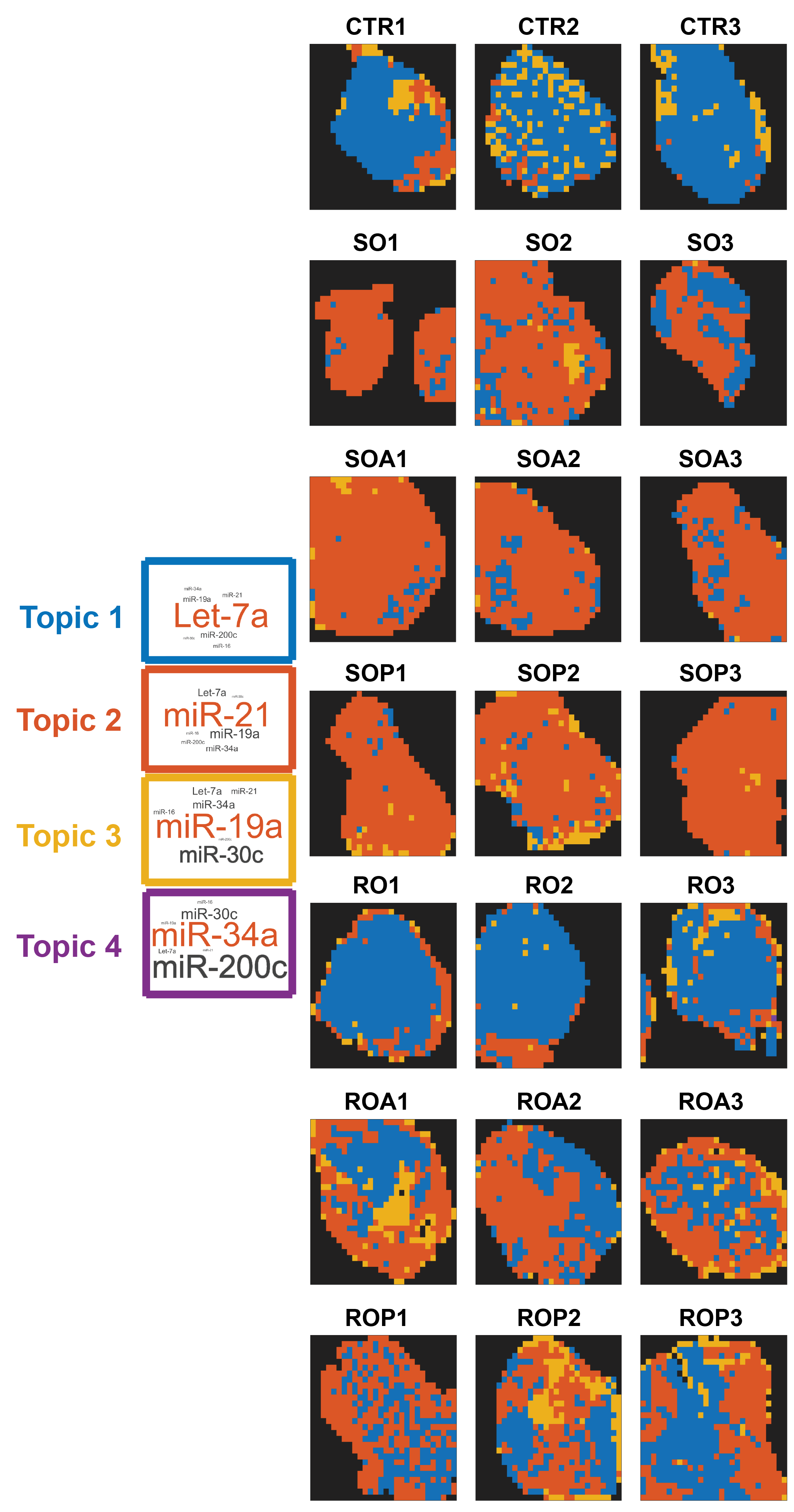

**S5 Fig. Case-study-wide spatial LDA dataset**. The full case-study-wide spatial LDA topic maps for all 10-day tumor samples. The color of the tumors corresponds to the topic bearing the same color.

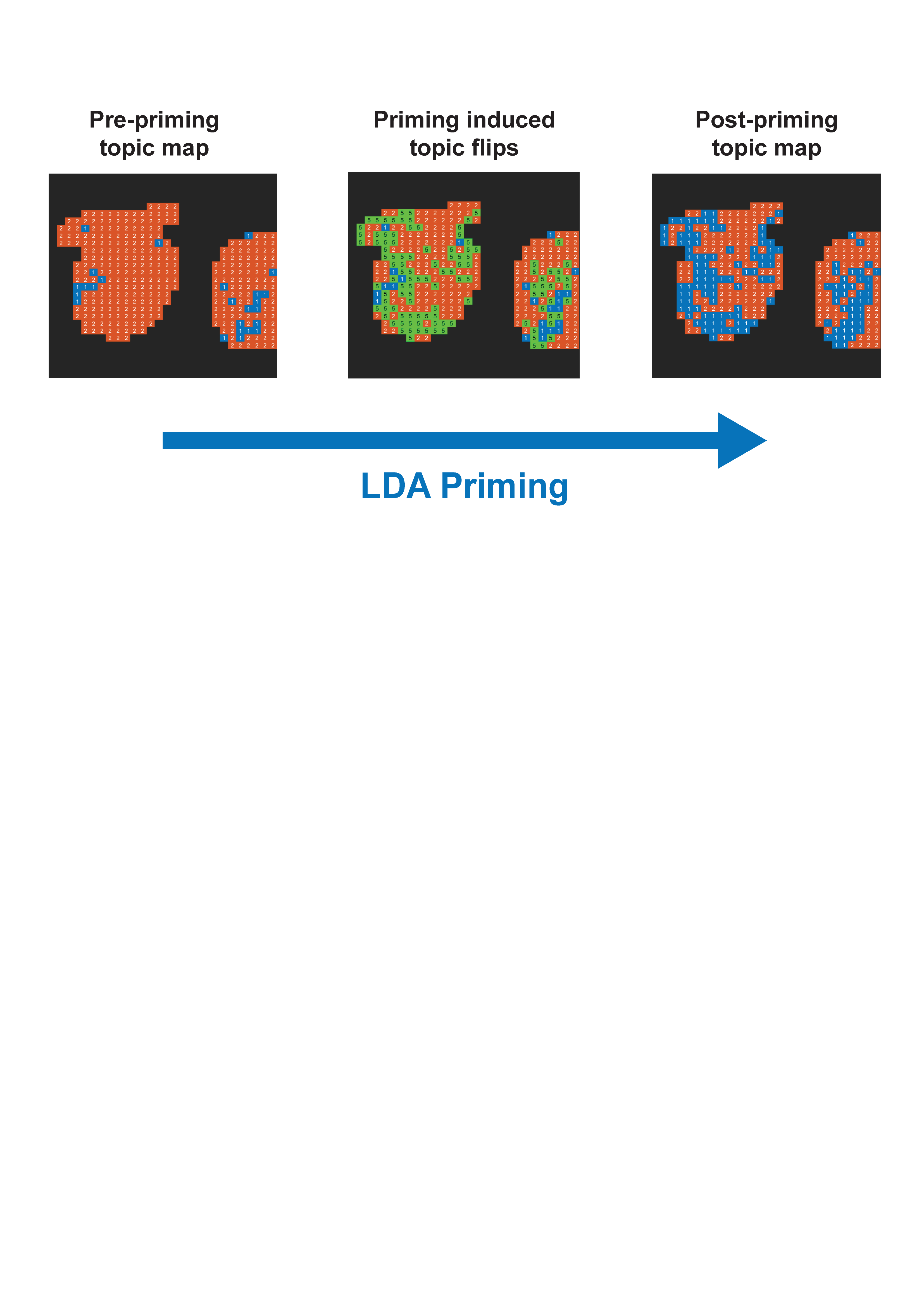

**S6 Fig. LDA topic changes induced by priming**. The application of LDA priming to the miRNA data results in changes (flips) of the determined dominant topic where the pre-biased miRNA amounts of let-7a and miR-21 were close in value. The green represents the locations dominant topic changed as a result of priming.

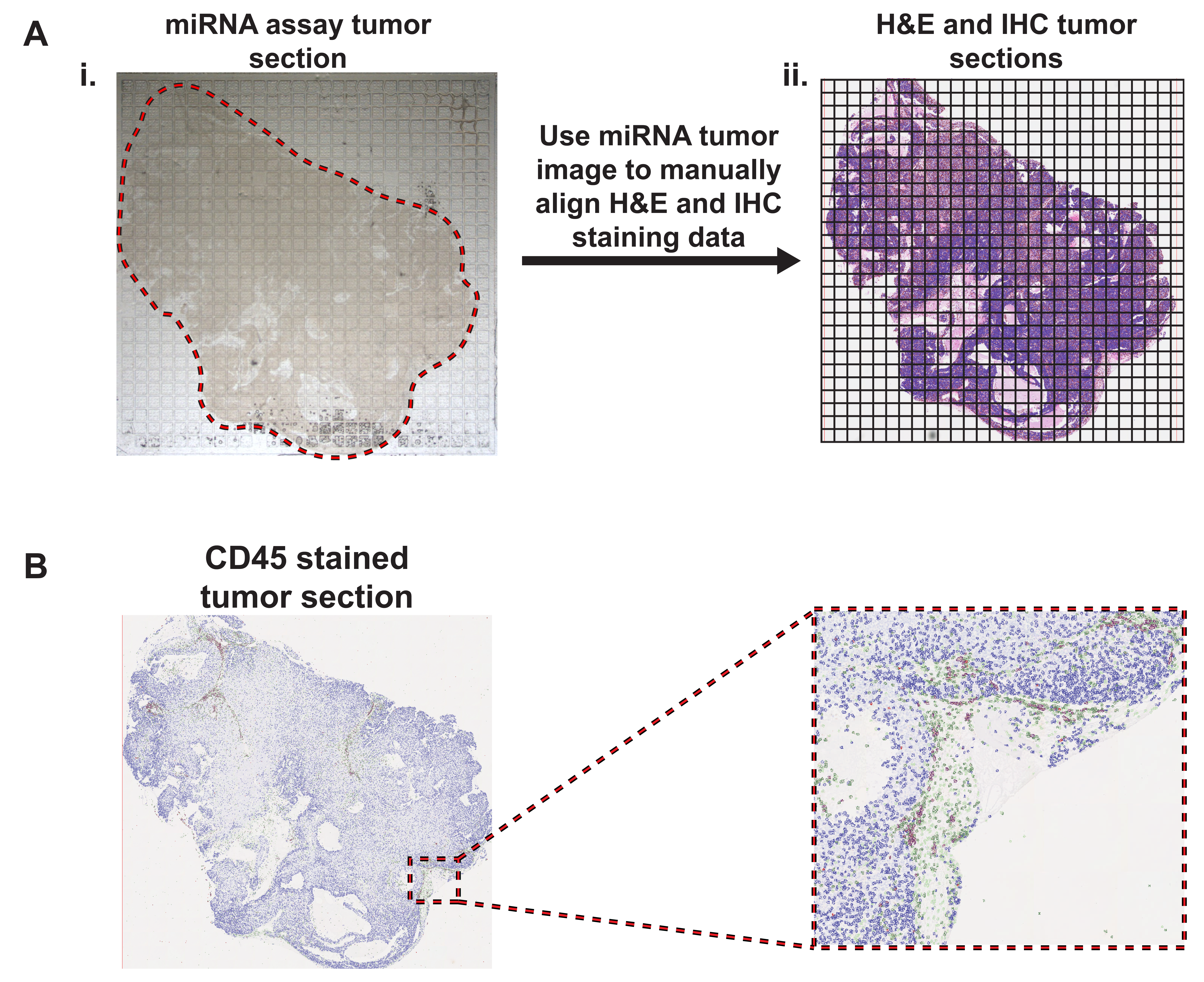

**S7 Fig. Tumor section alignment and CD45 staining**. **A. i.** Tumor sections are superimposed on the miRNA assay device and imaged before running the assay. The captured image is manually outlined to note where the tumor lies within the array. **ii.** A grid with the same dimensions as the assay device is then drawn on images of the adjacent tumor sections that are H&E and stained for CD45 such that they have approximately the same position within the array. This information is used downstream to calculate the number of cells within each well of the assay device via the QuPath derived data. **B.** High quality CD45 stained tumor sections were available for all the 10-day tumors.

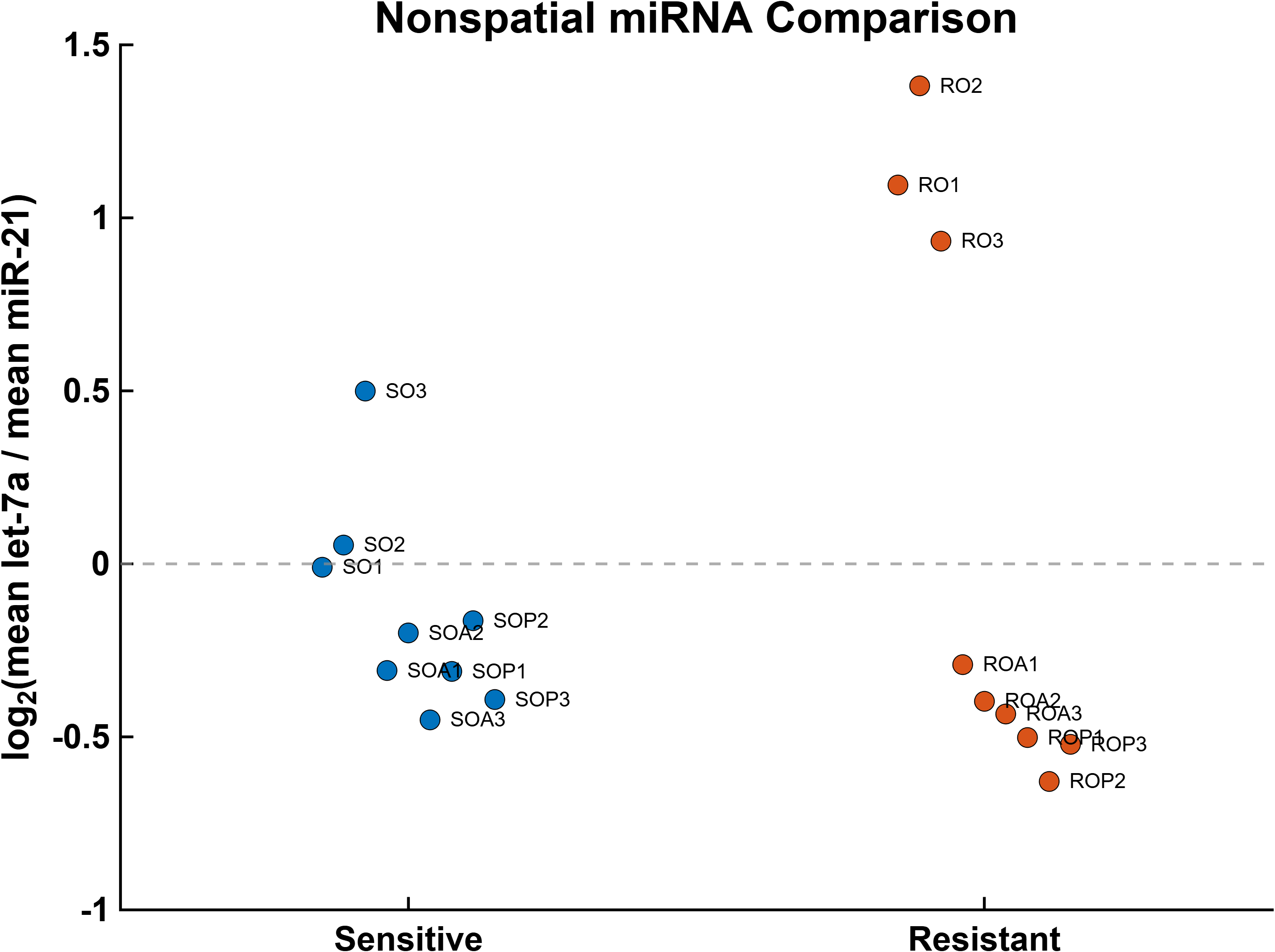

**S8 Fig. Nonspatial let-7a/miR-21 ratio comparison**. Tumor-level ratios of the mean let-7a to miR-21 measurements for olaparib-sensitive (n = 9) and olaparib-resistant (n = 9) tumors. Each point represents an individual tumor, and ratios are displayed on a log2 scale. No significant difference was detected between the two groups (exact two-sided Mann–Whitney U test, p = 0.546).
